# Non-caloric sweeteners combined with glucose affect hypothalamic glucose sensing-induced insulin secretion and refeeding through neuronal cellular metabolism: *in vivo* and *in vitro* approaches

**DOI:** 10.1101/2024.12.16.626024

**Authors:** J. Haydar, C. Fenech, F. Liénard, B. Abed, S. Grall, L. Briand, G. Walther, C. Leloup

## Abstract

Changes in brain activity associated with deleterious metabolic effects of non-caloric sweeteners (NCS) have been demonstrated in humans, particularly when their intake is concomitant with that of glucose. Here, we have focused on hypothalamic glucose sensing in rats, detecting increases in circulating glucose levels and in turn triggering various physiological controls. The identification of sweet-taste receptors in the hypothalamus has suggested that they participate in glucosensing mechanisms, but the existence of a dialogue between different pathways has never been studied. We tested the acute effects of hypothalamic glucosensing combined with an NCS (sucralose or acesulfame potassium (AceK)), the latter activating sweet-taste receptors without producing energy. Our working hypothesis was that the concomitance of two contradictory signals (energetic, sweet glucose vs. non-energetic, sweet NCS) could be responsible for deleterious physiological effects. After validation that sweet taste receptors and their signaling components were indeed present in the rat hypothalamus, insulin secretion induced by hypothalamic glucosensing (increased glucose level) in the presence of sucralose and AceK was examined. Insulin release was reduced compared to glucose alone, while the two NCS alone had no effect. Plasma norepinephrine, which had decreased after the glucose stimulus alone, instead rose when either NCS was co-injected with glucose, whereas the NCS had no effect with basal glucose. Regarding the satiety-inducing effect of glucose, concomitant injection of each NCS with glucose produced the opposite effect to that observed with glucose alone, with food intake being increased, an effect also observed with NCS injected alone. Using the GT1-7 hypothalamic cell line expressing sweet-taste receptors, we showed that the ATP concentration which normally increases with rising glucose levels was dose-dependently decreased in the presence of NCS, an effect no longer significant in the presence of gurmarin, an inhibitor of rodent sweet-taste receptors. The increase in ROS production in response to rising glucose levels was enhanced in the presence of NCS, an effect prevented by gurmarin. NCS also reduced substrate-stimulated mitochondrial respiration in permeabilized cells. Together, these results show that NCS combined with glucose disturb hypothalamic glucose sensing *in vivo* and impair mitochondrial energy metabolism in hypothalamic cells, through sweet-taste receptors.

## 2 | INTRODUCTION

Non-Caloric Sweeteners (NCS) first emerged as alternatives against metabolic disorders, promising sweetness without the caloric load [1,2]. However, recent studies in both humans and animals suggest that NCS might contribute to the development of type 2 diabetes and are positively correlated with weight gain [3,4,5,6]. Among the non-nutritive sweeteners approved for consumption by the US Food and Drug Administration (FDA) and by the European Food Safety Authority (EFSA), sucralose and acesulfame potassium (AceK) have risen in prominence, partly because of their significantly higher sweetness intensity compared to other NCS [7]. Sucralose and AceK are known to interact with the sweet taste receptor, a heterodimer of taste type 1 receptor 2 (T1R2) and taste type 1 receptor 3 (T1R3), primarily located on the taste buds on the tongue [8]. In addition to their well-known presence on the taste buds, sweet taste receptors have been identified in several extra-oral tissues, including the gastrointestinal tract, pancreas, adipose tissue, testis and brain [8]. The discovery of sweet taste receptors in these extra-oral locations has suggested that they may play roles beyond taste perception [9]. For example, in pancreatic beta cells, steviol glycosides (natural non-caloric sweet-tasting molecules) potentiate the activity of the TRPM5 cation channel – a channel involved in signal transduction and expressed in sweet-sensitive taste cells – and enhance glucose-induced insulin secretion in a TRPM5-dependent manner [10], demonstrating cross-talk between the sweet-taste receptor and classical glucose-induced insulin secretion (GIIS) metabolic signaling pathways.

Within the brain, sweet taste receptors have been identified in regions such as the hypothalamus and the reward-related circuits [11,12,13]. These areas are involved in regulating satiety, energy balance, and reward pathways [14]. In particular, the hypothalamus plays a central role in glucose sensing and metabolic regulation [15,16] through specialized neurons known as glucose-sensing neurons that are finely tuned to detect changes in blood glucose levels. In turn, they orchestrate appropriate physiological responses to maintain blood glucose level via changes in the activity of the autonomic nervous system that is essential for controlling carbohydrate metabolism [17,18]. For example, increased hypothalamic glucose levels stimulate the vagal control of insulin secretion that contributes to the first phase of glucose-induced insulin secretion [19]. These responses are driven by hypothalamic neurons sensitive to rise in blood glucose levels at least in the arcuate nucleus (ARC) [20,21,22,23]. The identity of these neurons is only partially known, however, due to the absence of a homogeneous neurochemical marker.

Until recently, this cascade of events was studied solely from an energetic perspective, demonstrating the crucial role of glucose oxidation and mitochondria-dependent signaling [24,20] and reviewed in [25,26]. It is only recently that the role of the sweet-taste receptor in hypothalamic glucose sensing has been investigated in ARC. To distinguish glucose sensing via the metabolic plus tasting receptor pathway from taste receptor glucose sensing alone, NCS sucralose has been used [27]. This NCS, from 10^−5^ to 10^−2^ M, increases intracellular calcium in a dose-dependent manner in 12-16% of freshly isolated ARC neurons, an effect that is abolished by the sweet taste receptor inhibitor gurmarin. These sucralose-responsive neurons are neurons otherwise activated by high concentrations of glucose (10 mM) whose response is attenuated in the presence of the sweet taste receptor inhibitor, gurmarin [27,28]. However, the acute, concomitant effect of elevated blood glucose with that of non-caloric sweeteners has never been explored at the hypothalamic level. This question is particularly important in view of a recent study showing that the concomitant consumption of sucralose with, but not without, carbohydrates reduced insulin sensitivity in humans, an effect correlated with a reduction in the activity of some brain areas, measured by fMRI [5]. However, when carbohydrates were consumed orally in combination with a non-caloric sweetener (sucralose) in humans, it was unclear why a defect in carbohydrate metabolism was observed. This could be due to sucralose binding to intestinal cells expressing the sweet taste receptor, thereby altering glucose absorption [29]. Another explanation could be a change in the central regulation of glucose metabolism, or both. In this study, we hypothesized that the response of hypothalamic glucose-responsive neurons to the simultaneous presence of glucose and a non-caloric sweetener affected at least the central, autonomic control of insulin secretion.

To test this hypothesis, we employed carotid injection to directly target the brain. This physiological approach has previously enabled us to demonstrate *in vivo* that the electrophysiological response of glucose-responsive ARC neurons plays a role in the nervous control of insulin secretion [20,21]. This method allowed us to compare the effects of increasing concentrations of glucose alone or non-caloric sweeteners alone (specifically, sucralose or acesulfame potassium (AceK)) with the combined effects of glucose plus a non-caloric sweetener. Additionally, we extended this study to evaluate the postprandial brain satiety signals typically induced by rising blood glucose levels by measuring refeeding after fasting in response to intraperitoneal injections of glucose alone, sucralose alone, AceK alone, or a combination of glucose plus sweeteners. Finally, to dissect how these different combinations might produce distinct effects at the cellular and molecular levels, we utilized the GT1-7 hypothalamic cell line. This cell line faithfully reproduces central glucose sensing which involves, among other markers, proteins associated with glucose sensing in pancreatic beta cells, such as the ATP-sensitive potassium channel (K_ATP_) [30,31,32,33,34,35]. Glucose-excited neurons of this cell line are inhibited when glucose concentration decreases (by increased opening of K_ATP_ channels, leading to neuronal hyperpolarization and firing inhibition) and are required *in vivo* to initiate appropriate counter-regulation to hypoglycemia [32,33].

We therefore used this cell line to investigate the mechanisms that might be involved in the uncoupling observed when the organism is specifically exposed to an increase in blood glucose in the presence of NCS, compared to an increase in blood glucose alone or sweeteners alone. The concurrent detection of glucose and non-caloric sweetening molecules could be at the origin of signaling perturbations involving the energy metabolism of glucose-excited neurons, in particular their ATP production. To test this possibility, we also conducted bioenergetics experiments. To explore this hypothesis, we conducted a series of bioenergetics experiments using the GT1-7 cell line, known for its glucose-sensing capabilities. These experiments aimed to determine if the concurrent presence of glucose and NCS could lead to disruptions in cellular energy metabolism. Specifically, we assessed mitochondrial respiration, ATP production, and ROS (reactive oxygen species) generation. By measuring these key indicators, we sought to understand how the combination of glucose and NCS might alter the normal functioning of glucose-excited neurons, potentially leading to metabolic uncoupling. Furthermore, we aimed to investigate whether these disruptions in cellular energy metabolism could, at least partially, explain the observed effects on insulin secretion during glucose sensing. We hypothesized that the activation of sweet taste receptors by NCS in the presence of glucose could alter the signaling pathways that regulate energy metabolism, leading to changes in ATP production and mitochondrial function. By focusing on whether these sweeteners exert their impact through a receptor-dependent mechanism, we sought to determine if the sweet taste receptors are integral to the metabolic changes observed. This dual activation by both glucose and sweeteners may disrupt normal cellular processes, contributing to the observed effects on insulin secretion during glucose sensing. Understanding this interaction could provide crucial insights into the mechanisms by which sweeteners influence central glucose metabolism and insulin regulation, highlighting the potential role of sweet taste receptors in these processes.

## 3 | MATERIALS AND METHODS

### 3.1. Animals and Housing

Male Wistar rats (7-8 weeks old, Charles River, France) were pair-housed as recommended for animal welfare and maintained in a temperature-controlled environment under a 12 h light/dark cycle (08:00 to 20:00), at 22°C and 40%-70% relative humidity in a conventional animal husbandry. Standard chow and water were provided ad libitum (#A04, Safe, France), except for groups whose food was removed 12 h before the refeeding test.

Surgeries and experiments were performed under acepromazine tranquilization (Calmivet, 0.25 mg/kg), alfaxalone for anesthesia (Alfaxan, 80 mg/kg, Centravet) and buprenorphine for analgesia (0.05 mg/kg, Buprecare). This study was approved by the local ethics committee of the University of Burgundy Europe (C2EA Grand Campus Dijon No. 105) and authorised by the French Ministry of Research (authorisation APAFIS #34709-2022030308579236 v4). All procedures complied with Directive 2010/63/EU of the European Parliament and of the Council on the protection of animals used for scientific purposes.

The sucralose, acesulfame potassium (AceK) and D-(+)-Glucose used in vivo and substrates used for mitochondrial respiration on GT1-7 hypothalamic cell line were purchased from Sigma (Sigma-Aldrich, St-Quentin-Fallavier, France).

### 3.2. RNA isolation and quantitative RT-PCR analysis

Anesthetized rats were killed by decapitation and their brains harvested immediately. Six brain regions were collected and rapidly frozen at –80°C: Hindbrain (HB), cerebellum (C), cortex (CX), hypothalamus (HT), thalamus (T), hippocampus (H), pancreas and tongue.

Total RNA from each brain region was extracted (Macherey-Nagel – NucleoSpin kit). RNA quality and quantity were assessed by microelectrophoresis (Experion, Bio-Rad or TapeStation, Agilent); samples with a RIN/RQI below 7 were not included in the study. Reverse transcription (PrimeScript™ RT, TaKaRa) was performed on standardized amounts of starting RNA for all samples.

Quantitative real-time PCR was carried out on a StepOnePlus thermocycler (Applied Biosystems) in duplicate, using hydrolysis probes (TaqMan Gene Expression Assays, Applied Biosystems) on standardized cDNA amounts (input RNA equivalent 70–100 ng). TaqMan assays for *Tas1r3, Tas1r2*, *Gnat3, Tas2r116, Gna14* and *Trpm5* are detailed in **Supplementary Table 1**; they were selected according to [36]. Genomic DNA removal steps are included during RNA extraction and before reverse transcription. Their efficiency is controlled with “no-RT” PCR reactions. Relative amounts of target cDNA were calculated by the comparative threshold (Ct) method (2−ΔΔCt) [37] using the qRAT package [38] with *Ppia* and *Pgk1* (rat samples) or *Ubc* and *Tbp* (mouse samples) as reference genes to normalize cDNA input. Both rat reference genes were found to be the most stably expressed across various regions of rat brain [39].

### 3.3. Experimental Groups

#### 3.3.1. Intra-carotid glucose injection-induced insulin secretion

Injection through the carotid artery (i.c.) was chosen as the most physiological route to the brain, since it is the route by which blood glucose and circulating molecules reach it. Our previous studies have demonstrated that hypothalamic glucose-excited arcuate (ARC) neurons are involved in the vagal control of the insulin peak induced by this intra-carotid glucose injection [20;21;23].

Two main groups were formed to study the effects of the 2 sweeteners on hypothalamic glucose sensing inducing insulin peak: a control group (“BG”) receiving a basal concentration of glucose (5 mM), non-stimulating in terms of hypothalamic glucose sensing (no vagal control triggered for insulin secretion) and a group receiving a stimulating concentration of glucose-provoking cerebral hyperglycemia named “HG” (9 mg/kg) [23;40]. In both groups, the effects of sweeteners were tested: sucralose (1 mM) and AceK (3 mM) [41], groups named Sucra+BG and AceK+BG in the basal group, respectively; and groups named Sucra+HG and AceK+HG in the “HG” group, respectively. Each group consisted of eight rats and surgeries were performed as previously described on anesthetized rats [23;40]: a polyethylene catheter was inserted into the carotid artery and advanced 5 mm in the cranial direction, then ligatured. A bolus of the different solutions tested was injected in 100 µL of saline toward the brain over 30 s, at a flow rate of 200 µL/min. Blood glucose was measured from a drop of tail-vein blood before (T0) and at 1 (peak insulin), 3, 5 and 10 min after the load. Blood for plasma insulin was collected from the tail vein into Microvette® tubes (Sarstedt) at the same times. Additional blood was collected in separate tubes at T0, 1 and 10 min for norepinephrine. Samples were centrifuged at 7000 rpm for 10 min at 4°C, and plasma was stored at −20°C until analysis.

#### 3.3.2. Blood parameters determination

Blood glucose concentration was measured at the indicated times using the glucose analyzer Performa AccuCheck (Roche Diagnostics, France). Plasma insulin (Insulin ALPCO kit, Eurobio, France) and norepinephrine (Cusabio, CliniSciences, France) concentrations were measured using ELISA kits according to the manufacturer’s instructions. Values are expressed in ng/mL.

#### 3.3.3. Refeeding test

After a 1-week acclimatization period, rats (250–300 g) were randomly assigned to six groups (n = 8 per group). After an overnight 12-h fast, rats received an i.p. injection of glucose alone (2 g/kg; positive control [42]), sucralose alone (20 mg/kg [43]), AceK alone (100 mg/kg [44]), sucralose + glucose or AceK + glucose (same doses). Control rats (negative control) received PBS. The same volume was injected in all groups. Food was returned 15 min after the injection, and food consumption was measured at 30, 60, 120, 180 and 240 min after injection and normalized to body weight.

### 3.4. An in vitro model to assess effects of sweeteners

#### 3.4.1. Expression of taste receptors in GT1-7 cell line

To compare levels of taste receptors (*Tas1r3*, *Tas1r2*) and signaling protein (*Gnat3*) transcripts in mouse hypothalamus and hypothalamic cell line GT1-7, RT-qPCR was carried out on total RNA from three equivalent cultures of GT1-7 cells as well as RNA isolated from the hypothalami of 5 mice, as described above. Expression levels were normalized to the mean of *Ubc* and *Tbp* expression levels. Transcripts and corresponding TaqMan® ID (Thermo Fisher Scientific, Waltham, MA, USA) are provided in full in **Supplementary Table 1**.

#### 3.4.2. Effects of sweeteners on ATP level and mitochondrial respiration of GT1-7 hypothalamic cell line exposed to glucose

The GT1-7 hypothalamic cell line (Merck, France), a model for the study of hypothalamic glucose sensing in vitro [32,33] was cultured within Dulbecco’s Modified Eagle’s Medium (DMEM high-glucose medium-25 mM glucose) containing 10% fetal bovine serum (FBS, Cytiva, MA, USA) and supplemented with 1% pen/strep (Gibco). Cells were maintained at 37 °C and 5% CO2 conditions. Cell viability was assessed by trypan blue exclusion before each experiment, and only suspensions with over 85% viable cells were used for respiration experiments (98% for the ROS assays). To rule out toxicity of the test compounds, viability was also measured after 30 min of exposure to the highest concentrations of sucralose (5 mM) and AceK (15 mM); it did not differ from that of the glucose controls (**Supplementary Fig. S1**).

***ATP determination***. To assess ATP levels in GT1-7 cells exposed to NCS, 120 × 10^3^ cells (passage up to 16) were seeded in 12-well plates in low glucose medium (2.5 mM) at least 48 hours before use. Sucralose (0.01 mM, 0.1 mM, 1 mM and 5 mM) or AceK (0.03 mM, 0.3 mM, 3 mM and 15 mM) was added to the culture medium for dose-response experiments for 30 min. Sucralose and AceK concentrations were in accordance with concentrations used by other authors for in vitro experiments [27;45]. Cells were stimulated with 20 mM glucose, with or without NCS at different concentrations, and with or without the sweet taste-suppressing peptide recombinant Q1-gurmarin (10 µg/ml, provided by L. Briand, Dijon, France; details in **Supplementary Appendix**).

After incubation, the medium was rapidly removed, and phenol-chloroform was immediately added to the cells thus inactivating ATPases [46]. After phenol/chloroform/water extraction the ATP/DNA-containing aqueous phase was collected and used directly for ATP concentration determination. ATP concentration was measured on 20 µL aqueous phase using a luciferin/luciferase bioluminescent assay (Invitrogen). Genomic DNA concentration in the aqueous phase was also measured for all samples to normalize ATP amounts to cell amounts in each well. Wells only treated with 20 mM glucose were used as positive controls. Experiments were repeated at least four times each with duplicate wells.

ROS quantitation/Generation. For the quantification of reactive oxygen species (ROS) in GT1-7 cells, the Amplex® Red Hydrogen Peroxide/Peroxidase Assay Kit (Invitrogen) was employed. This assay was conducted under stringent conditions to ensure the accuracy and reproducibility of results. All buffers were pre-warmed to 37°C, and the Amplex Red reagents were protected from light exposure throughout the experiment to prevent degradation. **Cell Culture and Preparation**: GT1-7 cells were cultured in a low glucose medium for at least 48 hours before the assay. On the assay day, cells were harvested and resuspended in the low glucose medium. Cell counting and viability assessment were conducted, ensuring a high viability rate of 98%. **Experimental Setup**: For each experimental condition, 100 µL of the reaction mixture was dispensed into the wells of a 96-well plate. The experimental conditions included: Low glucose (2.5 mM), High glucose (20 mM), High glucose (20 mM) NCS (Sucralose or AceK), both at 1 mM, with or without the addition of gurmarin (10 µg/ml), rotenone was used as a positive control for inducing the production of hydrogen peroxide (H₂O₂) in the cell. Following the addition of 20 µL of cell suspension (at a concentration of 9.34 x 10^^5^ cells/mL) to each well, the plate was incubated for 30 min to allow the cells to acclimate and react to the different conditions. Fluorescence readings were taken every five min over a 60-min period to track ROS production. Amplex Red detects extracellular H₂O₂; throughout this article, ‘ROS generation’ refers to this measurement. **Data Collection and Analysis**: Fluorescence was measured using an excitation range of 530–560 nm and emission detection at approximately 590 nm. Alternatively, absorbance was measured at approximately 560 nm. Data are reported as the delta value, which represents the difference between readings at time = 0 and at each 5-min interval. Readings were recorded for 60 min. Only the first 20 min are shown, because no differences between conditions were observed after 20 min

***Mitochondrial respiration***. Oxygen consumption was measured using a respirometer (Oxygraph-2k; Oroboros Instruments). Between 2 and 3 million GT1-7 cells were transferred into mitochondrial respiration medium (MiR05) containing saponin for permeabilization (2.5 µL of freshly prepared saponin solution (5 mg/ml MIR05 buffer)), according to previous protocol [47]. Then, cells were transferred into the glass chamber (1.8 ml) of the respirometer. The baseline of the respiration was first measured, then mitochondrial state 2 respiration was stimulated by the addition of 20 mM succinate (called substrate-driven respiration) before or after increasing concentrations of sweeteners (10^−3^, 10^−2^ or 1 mM of sucralose or AceK) to measure its impact according to the mitochondrial status (non-stimulated vs. stimulated). Finally, oxygen consumption per million of cells and per second was calculated using DatLab software. Media were prepared according to the guide provided by Oroboros Instruments, with technical sheets available on the company website at www.oroboros.at.

### 3.5. Statistical analysis

All data were expressed as mean ± standard error of mean (S.E.M.). Statistical analysis was performed using GraphPad Prism Software (version 10.03, GraphPad Software, San Diego, CA, USA). One-way/ two-way analysis of variance (ANOVA) followed by the Bonferroni, Dunnett’s or Tukey post hoc tests was used for analysis of differences between control and experimental groups or paired analyses was used for analysis of differences between control and experimental groups. Values of p < 0.05 indicate statistical significance between the compared groups. For PCR analyses, values are presented as relative quantity (RQ) ± standard error of the mean (SEM), derived from a minimum of two biological replicates, and normalized to the reference genes, unless stated otherwise. Comparison of two treatments was done using Student’s t-test for normally distributed data and the Mann-Whitney U test for non-normally distributed data. whereas one-way ANOVA followed by post hoc test was applied for comparison of multiple treatments.

## 4 | RESULTS

### 4.1. Quantification of sweet and bitter taste receptor mRNAs and key downstream signaling components across rat brain regions

Real-time PCR for *Tas1r2*, *Tas1r3*, *Gnat3*, *Trpm5*, *Gna14* and bitter taste receptor *Tas2r116* was used to quantify the expression profiles of these genes in Wistar rats throughout different brain areas including the hypothalamus (**HT**), the cortex (**CX**), the hippocampus (**H**), the cerebellum (**C**), the thalamus (**T**) and the hindbrain (**HB**). Pancreas was chosen as a well-documented glucosensing organ through T1R2/T1R3 pathway, and tongue, as a reference organ for sweet taste [48]. We compared the mRNA levels of these genes with those of the hypothalamus, taken as 100% reference, due to its role in hypothalamic glucose sensing. First, we analyzed mRNA expression of the subunit *Tas1r2* (**Fig. 1A**). *Tas1r2* mRNA was more highly expressed in the hypothalamus than all other brain areas (p<0.0001), the thalamus being the second brain area with a 20% *Tas1r2* mRNA expression. *Tas1r2* mRNA was almost undetected in the pancreas. It was, however, about three times less expressed in the hypothalamus than in the tongue (p<0.0001). Second, we quantified the mRNA of the other subunit in the heterodimeric sweet taste receptor, *Tas1r3*. Cerebellum presented the highest expression among brain areas (250%), i.e., 2.5 times higher than the HT (100%) (p<0.001) (**Fig. 1B**). Unlike *Tas1r2* mRNA level in the pancreas, *Tas1r3* mRNA showed a substantial expression compared to the hypothalamus. Both tongue (640%) and pancreas (378%) expressed higher levels of *Tas1r3* mRNA compared to hypothalamus (p<0.0001). We next analyzed the Gα-gustducin (*Gnat3*) mRNA levels, a specific pathway for transducing signals through some taste receptor [49] (**Fig. 1C**). Comparison between brain areas only shows that *Gnat3* mRNA levels are more highly expressed in the HT than all other brain areas (p<0.0001, except thalamus, p<0.002). However, the largest expression was found in pancreas (3620%) and tongue (3850%) (p<0.0001 compared to each brain area). With regard to the expression profile of *Trpm5*, a sodium-selective TRP channel that acts as a common downstream component to transduce taste stimuli into output signals [50], the profile was similar to that of *Gnat3* (**Fig. 1D**) with *Trpm5* mRNAs expressed in all brain areas, but displaying 5– and 7-fold lower levels in the hypothalamus compared with the tongue and pancreas, respectively (p<0.0001). The G-protein α subunit Gα14 also participates in the taste signal transduction [51,52], and its mRNAs were quantified. As shown in **Fig. 1E**, *Gna14* mRNA levels were significantly higher in the tongue and pancreas compared to the hypothalamus, at 471% and 310% of hypothalamic levels, respectively (p < 0.0001). In the brain, both hippocampus and cerebellum exhibit 2 times higher expression than the hypothalamus (206.7% ± 33.4% and 218.3% ± 32.1% vs. 100% ± 7.2%, p<0.05 and p<0.01, respectively). Finally, we analyzed the mRNA levels of the bitter taste receptor *Tas2r116*, also identified in brain [53]. *Tas2r116* mRNA levels were barely detectable in the pancreas and thalamus, at 26% and 14% of hypothalamic levels, respectively. The highest brain expression was observed in the hippocampus, at 235% of hypothalamic levels (p < 0.01). In contrast, the hypothalamus exhibited mRNA levels nearly 7-fold lower than those in the tongue (735%, p < 0.0001). This brain mapping and quantification of taste receptor mRNAs and associated signaling reveal distinct expression profiles for sweet taste subunits: *Tas1r2* mRNAs predominate in the hypothalamus, while *Tas1r3* mRNAs are more prevalent in the cortex, hippocampus, and cerebellum.

**Figure 1:**
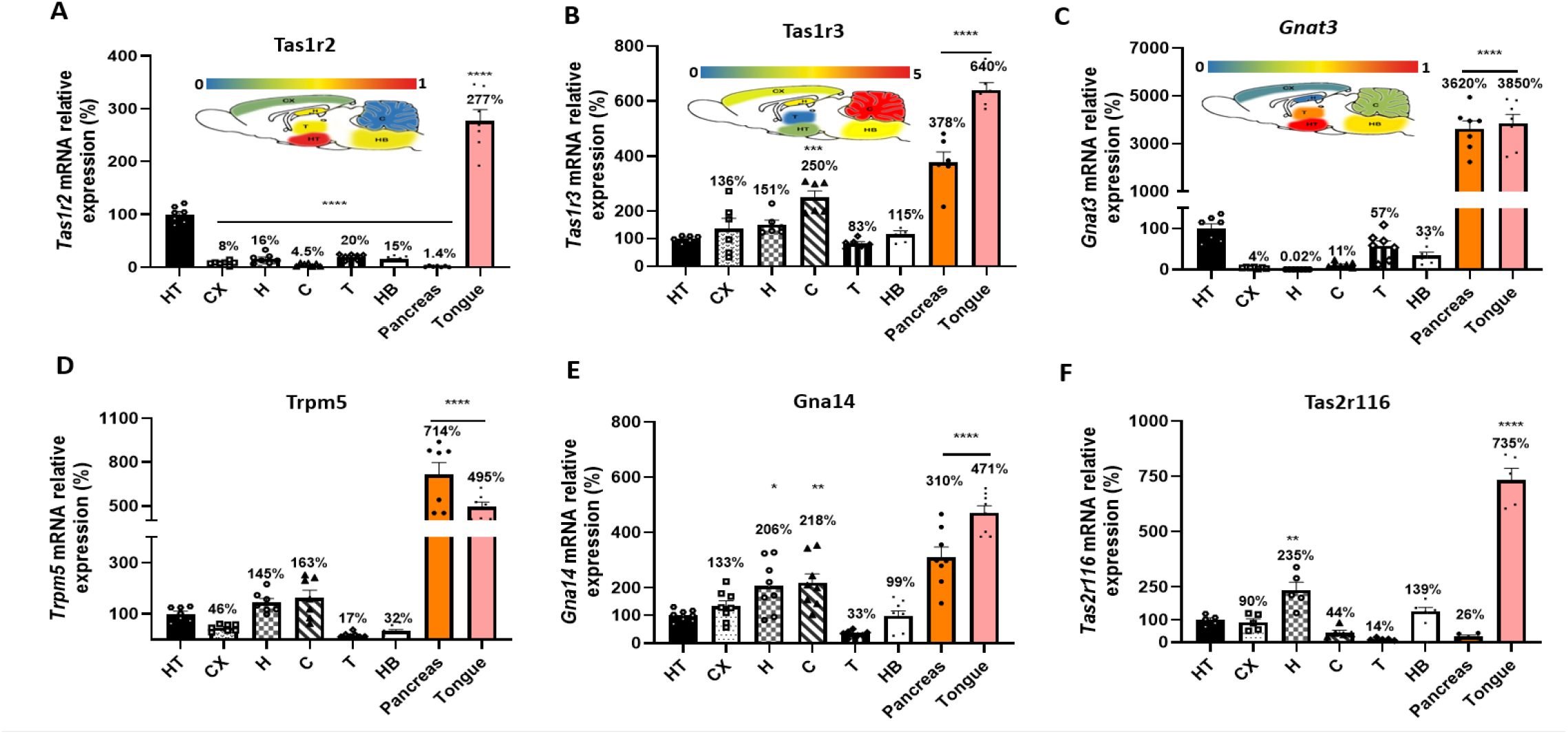
Relative mRNA expression of Sweet (*Tas1r2* and *Tas1r3*) and Bitter (*Tas2r116*) Taste Receptors in major brain areas and downstream signaling pathways compared to tongue (reference organ) and pancreas in Wistar rat. Relative Quantity (RQ) values are expressed as percentages, with the hypothalamus used as the reference point for comparison and set to 100% (**A**) Sweet receptor subunit *Tas1r2* mRNA expression in different brain areas (hypothalamus (HT), the cortex (CX), the hippocampus (H), the cerebellum (C), the thalamus (T) and hindbrain (HB)), pancreas and tongue as assessed by real time RT-qPCR. (**B**) Sweet receptor subunit *Tas1r3* mRNA expression in different brain areas compared to the pancreas and tongue. (**C**) Expression of Gα-gustducin (*Gnat3*) in different brain areas compared to the pancreas and tongue. (**D**) *Trpm5* mRNA expression in different brain areas. Cerebellum presents the highest expression in brain. *Trpm5* is highly expressed in the pancreas. (**E**) *Gna14* mRNA expression in the brain. Higher brain expression levels were found in the hippocampus and cerebellum. *Gna14* is highly expressed in the pancreas. (**F**) Bitter receptor *Tas2r116* mRNA expression in brain areas. The hippocampus presents the highest expression level in the brain. *Tas2r116* is barely detected in the pancreas. Schematic diagrams of the rat brain atlas are shown for *Tas1r2*, *Tas1r3*, and *Gnat3* expression levels. Color coding indicates expression levels in each brain region, with the color scale reflecting the range of RQ means: brighter blue represents lower expression (0), and darker red represents higher expression (1 or 5). mRNA expression was normalized to *Pgk1* and *Ppia*. Statistical comparisons between tongue, pancreas, and different brain areas versus the hypothalamus were performed using one-way ANOVA. *p < 0.05; **p < 0.01; ***p < 0.001. Values are presented as the mean ± standard error of the mean (n = 5-8).

### 4.2. Intra-carotid loading of sucralose and AceK with, but not without, stimulating glucose, impaired hypothalamic glucose-induced insulin secretion

The hypothalamo-pancreatic axis was stimulated by injecting a 30 s carotid load of basal or stimulant glucose, with or without sweeteners, into the brain. By this route, peripheral blood glucose levels are unaffected by the glucose load (low or high dose). A stimulating dose of glucose activates hypothalamic glucose-excited neurons [20;21;22]. This activation triggers a peak in insulin secretion through pancreatic vagal nerve activity within 1-3 min [23]. The experimental design and the conditions tested are described in **Fig. 2A**. Under normal conditions, the 30 s high glucose (HG) load into the carotid artery toward the brain caused a rapid peak of plasma insulin 1 min after the carotid injection in normal rats (T0 min vs. T1 min, delta plasma insulin = +2.36 ± 0.09 ng/ml, p < 0.0001) (Fig. 2B). The basal glucose (BG) load was ineffective to trigger a similar insulin peak (**Fig. 2B**), whereas blood glucose levels remained unchanged (see inset in **Fig. 2B**).

**Figure 2:**
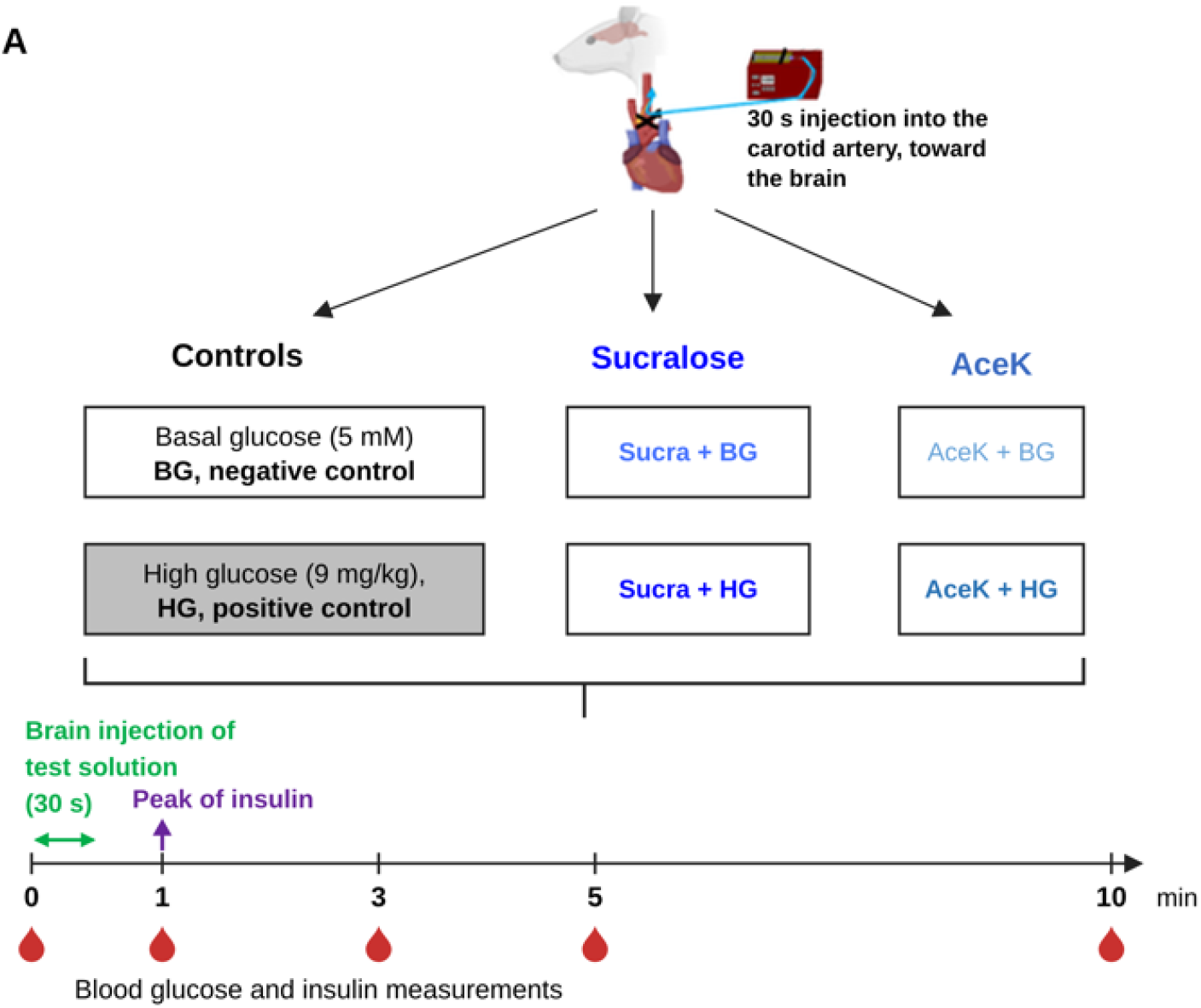
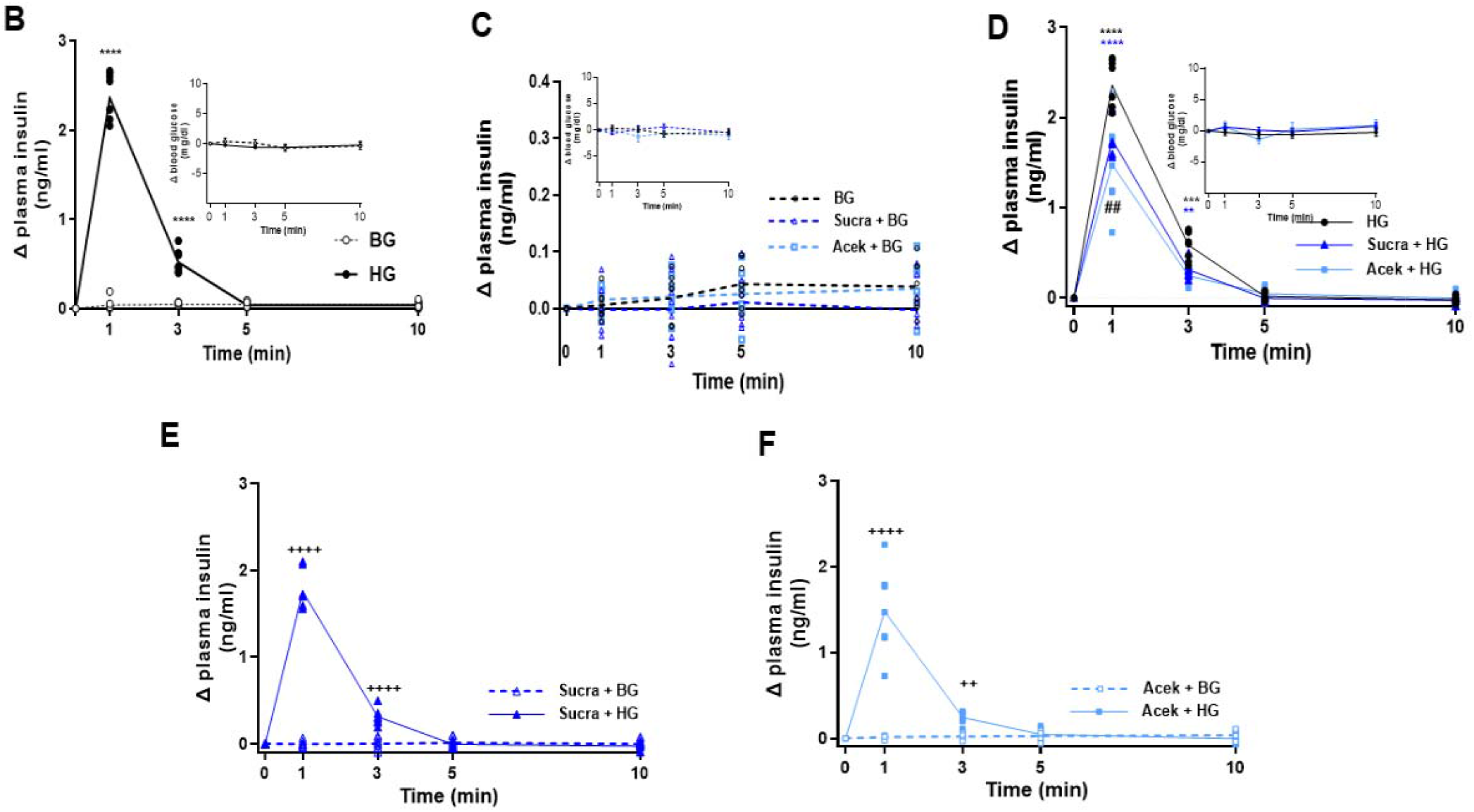
Rats display impaired hypothalamic glucose sensing-induced insulin secretion after a simultaneous intra-carotid injection of non-caloric sweeteners (NCS) and glucose. (**A**) Experimental procedure of the hypothalamic glucose sensing test with or without NCS. Basal glucose (BG) mimics the basal blood glucose concentration (i.e., no change), whereas the 9 mg/kg glucose load (HG, high glucose) is the dose known to stimulate glucose-excited neurons and trigger an insulin peak without altering peripheral blood glucose (indicating vagal control originating in the hypothalamus). Six groups were established: negative control (BG, no insulin peak), positive control (HG, insulin peak at 1 min), NCS alone (sucralose or acesulfame K [AceK] with BG), and NCS with glucose (sucralose or AceK + HG). In all conditions, the solution was injected through the carotid artery toward the brain for 30 s. (**B**) Plasma insulin in the normal physiological state, expressed as the difference from time 0 (before injection) to assess the peak, after intra-carotid injection of HG compared with BG. (**C**) Plasma insulin, expressed as the difference from time 0, after intra-carotid injection of BG (5 mM glucose), Sucra+BG (5 mM glucose + 1 mM sucralose) or AceK+BG (5 mM glucose + 3 mM AceK). (**D**) Plasma insulin, expressed as the difference from time 0, after intra-carotid injection of HG (9 mg/kg glucose), Sucra+HG (9 mg/kg glucose + 1 mM sucralose) or AceK+HG (9 mg/kg glucose + 3 mM AceK). (**B–D**) The inset of each graph shows the change in blood glucose between T0 (before injection) and the subsequent times after injection in each group. No significant difference in blood glucose was observed under any injection condition. Graphs (**E**) and (**F**) are excerpts from graphs C and D, respectively, allowing a clearer side-by-side comparison of Sucra+BG and Sucra+HG (E), and of AceK+BG and AceK+HG (F). Results are mean ± SEM (n = 8 per group). Data were analyzed by two-way ANOVA followed by Bonferroni’s multiple comparisons test; p-values are adjusted for multiple comparisons. Asterisks indicate significant differences from the positive control group (HG): ****p < 0.0001 (HG vs Sucra+HG and vs AceK+HG at 1 min); **p < 0.01 (HG vs Sucra+HG at 3 min); ***p < 0.001 (HG vs AceK+HG at 3 min). ##p < 0.01, Sucra+HG vs AceK+HG at 1 min. ++p < 0.01 and ++++p < 0.0001, each sweetener with BG vs the same sweetener with HG (E, F; at 1 and 3 min)

To investigate whether sweeteners alone, without a stimulating concentration of glucose, could activate glucose-excited neurons and trigger an insulin peak, a combined solution of either sucralose or AceK with basal glucose (BG) was injected. Under these conditions, neither sucralose nor AceK induced an insulin peak when glucose concentration was at basal levels (**Fig. 2C**). There was no significant difference in insulin levels between time 0 (before injection) and subsequent time points in the “basal glucose” group, the “basal glucose + sucralose” group, or the “basal glucose + AceK” group (**Fig. 2C**). Additionally, blood glucose levels remained unchanged (see inset in **Fig. 2C**).

In contrast, when NCS were injected simultaneously with a stimulating concentration of glucose to induce an insulin peak, this peak was altered (**Fig. 2D**). As expected, the positive control group (“High glucose,” HG) showed an insulin peak one min after injection. However, this peak was significantly decreased in the two groups that received simultaneous sweetener and glucose injections compared to the glucose-only group (HG group: T0 vs. T1 min = +2.36 ± 0.09 ng/ml, p < 0.0001, and T0 vs. T3 min = +0.58 ± 0.05 ng/ml). In the “High Glucose + sucralose” group (Sucra+HG), the insulin increase was +1.75 ± 0.07 ng/mL at T1 min (p < 0.0001), with a significant difference between HG and Sucra+HG at 1 min (p < 0.0001) and 3 min (p = 0.003). Similarly, in the “High Glucose + AceK” group (AceK+HG), the insulin increase was +1.47 ± 0.16 ng/mL at T1 min (p < 0.0001), with a significant difference between HG and AceK+HG at 1 min (p < 0.0001) and 3 min (p < 0.001). Additionally, a significant difference was noted between the Sucra+HG and AceK+HG groups at T1 min (p = 0.003). No significant change in peripheral blood glucose levels was observed across any of the groups, as shown in the figure inset.

To further distinguish between the impact of NCS alone versus their combination with glucose, **Figs. 2E and 2F** compare (Sucra + BG) versus (Sucra + HG) and (AceK + BG) versus (AceK + HG) respectively. In these conditions, both NCS failed alone (with BG) to produce any significant impact on insulin levels. In contrast, when combined with a high glucose concentration (HG), these NCS led to insulin release, manifested as an insulin peak at T=1 min. These results indicate that NCS alone do not stimulate the hypothalamic mechanisms involved in insulin secretion. However, when sweeteners are combined with increased cerebral glucose levels, they appear to diminish the nervous control of insulin secretion, leading to altered insulin response.

### 4.3. Non-caloric sweeteners increase NE plasma concentration during acute cerebral hyperglycemia, in contrast to a decrease during cerebral hyperglycemia alone

In previous studies, including work by [23;54], we and others have demonstrated that insulin peaks involve vagus nerve activity. To investigate whether the sympathetic autonomic response during cerebral hyperglycemia might also be influenced by NCS, we measured plasma norepinephrine (NE) concentrations 10 min after a 30-second intra-carotid injection of glucose or sweeteners in rats. This analysis compared conditions with stimulating glucose concentrations (high glucose, HG) versus basal glucose (BG), with or without the addition of sucralose (Sucra+BG or Sucra+HG) or AceK (AceK+BG or AceK+HG), and we also measured NE levels before the injection at T=0.

Our results showed that cerebral hyperglycemia alone led to a significant decrease in plasma NE concentration, with NE levels dropping from 0.162 ± 0.002 ng/mL in the basal glucose (BG) group to 0.152 ± 0.002 ng/mL in the high glucose (HG) group (p<0.01) (**Fig. 3A**). This decrease was not observed in the absence of stimulating glucose, where the presence of non-caloric sweeteners did not alter NE levels significantly (**Fig. 3B**). When we repeated the protocol with combined exposure to stimulating glucose and non-caloric sweeteners, we observed a reversal of the NE decrease associated with hyperglycemia alone. Rats exposed to hyperglycemia in the presence of sucralose or AceK showed a significant increase in plasma NE concentrations compared to the HG group. Specifically, NE levels increased from 0.152 ± 0.002 ng/mL in the HG group to 0.182 ± 0.002 ng/mL in the Sucra+HG group (p<0.0001) and from 0.152 ± 0.002 ng/mL to 0.182 ± 0.001 ng/mL in the AceK+HG group (p<0.0001) (**Fig. 3C**). There was no significant difference between the sucralose and AceK groups in these conditions. We also measured plasma norepinephrine (NE) concentrations at 1 mi following the 30-second intra-carotid injection to correspond with the time of the insulin peak. This additional measurement was aimed at understanding the impact of glucose and non-caloric sweeteners on sympathetic activity during this critical period. In the control groups (**Fig. 3D**), where basal glucose (BG) was administered, NE levels were measured at both T=0 (before injection) and T=1 min (during the insulin peak). No significant difference in NE concentrations was observed between these two time points, indicating that basal glucose did not substantially affect NE levels during the insulin peak. For the hyperglycemia (HG) group, a notable decrease in NE levels was observed at T=1 min compared to T=0. Specifically, NE levels dropped from 0.160 ± 0.002 ng/mL at T=0 to 0.150 ± 0.003 ng/mL at T=1 min (p<0.05). Suggesting that acute hyperglycemia is associated with a reduction in sympathetic tone during the insulin peak. When the same measurements were repeated with non-caloric sweeteners, distinct patterns emerged. In the sucra+BG group (**Fig. 3E**), NE levels at T=1 min did not differ significantly from those at T=0, indicating that the addition of sucralose to basal glucose did not alter NE concentrations at the time of the insulin peak. In contrast, the sucra+HG group showed a significant increase in NE levels at T=1 min compared to T=0. NE levels increased significantly from 0.159 ± 0.002 ng/mL at T=0 to 0.181 ± 0.003 ng/mL at T=1 min (p<0.0001). This indicates that sucralose combined with hyperglycemia counteracts the typical decrease in NE levels associated with hyperglycemia alone (**Fig. 3E**).

**Figure 3:**
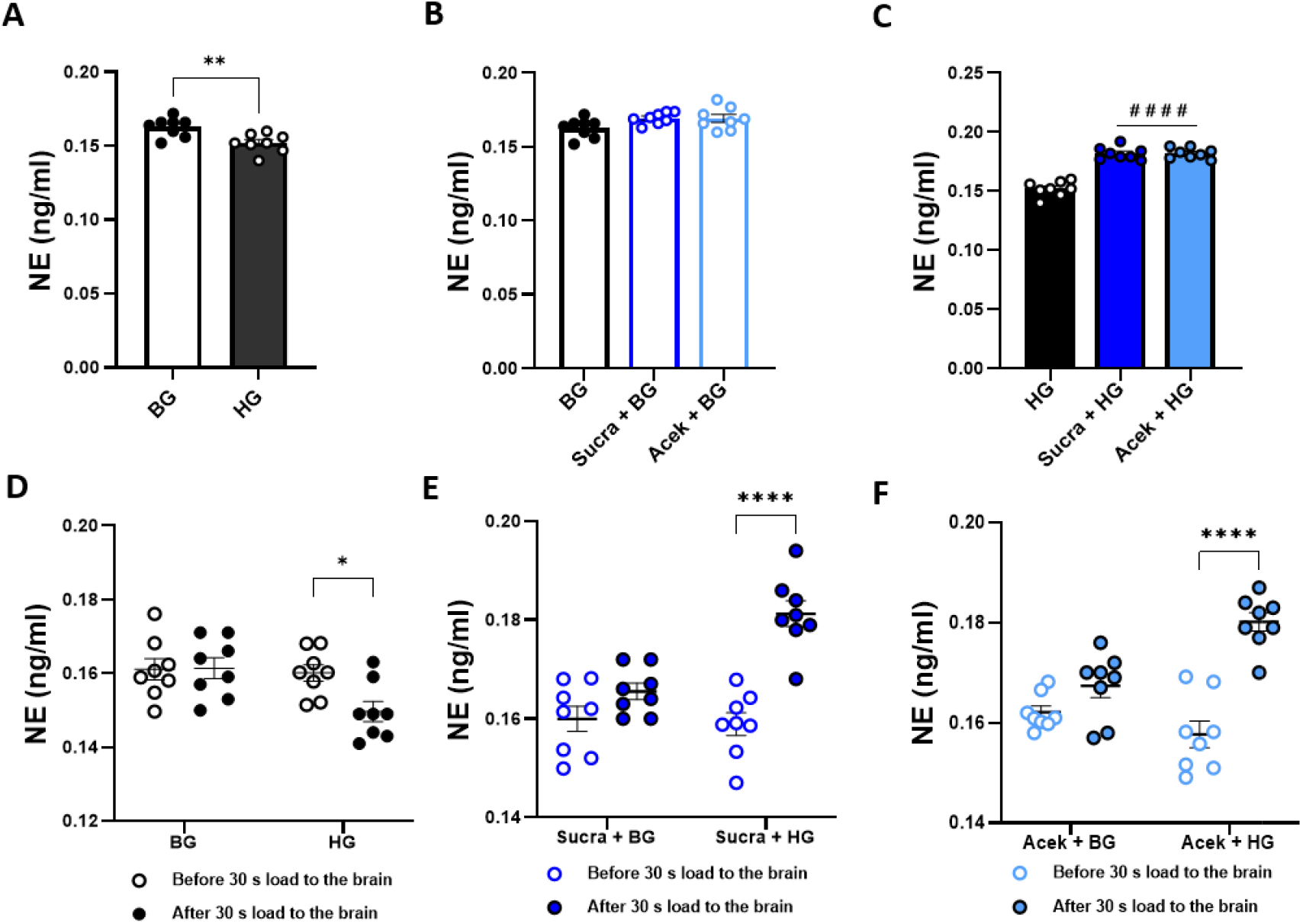
Non-caloric sweeteners increase plasma norepinephrine (NE) concentration after acute cerebral hyperglycemia, whereas cerebral hyperglycemia alone decreases it. (**A–C**) Plasma NE concentration 10 min after a 30-s intra-carotid injection of different solutions. (**A**) NE levels after a glucose solution mimicking hyperglycemia (9 mg/kg, HG group) vs a control group receiving basal glucose (BG, 5 mM). (**B**) NE concentrations after intra-carotid injection of NCS (sucralose or AceK) combined with BG, compared with BG alone. (**C**) Effect of combining NCS (sucralose or AceK) with HG, compared with HG alone. (**D–F**) Individual plasma NE concentrations before (T0) and after the 30-s injection. (**D**) BG and HG groups. (**E**) Sucralose combined with BG or HG. (**F**) AceK combined with BG or HG. Differences were assessed by one– and two-way ANOVA followed by Bonferroni’s post hoc test. Results are mean ± SEM (n = 8 per group; assays in duplicate). *, HG compared with BG (with or without NCS); #, NCS (sucralose or AceK) combined with HG compared with HG alone.

A similar trend was observed in the AceK group, where NE levels at T=1 min were significantly higher than at T=0, indicating that AceK also mitigates the decrease in NE levels associated with hyperglycemia. NE levels rose significantly from 0.158 ± 0.003 ng/mL at T=0 to 0.180 ± 0.002 ng/mL at T=1 min (p<0.0001). This finding shows that AceK also mitigates the reduction in NE levels seen with hyperglycemia (**Fig. 3F**). These results suggest that NCS, specifically sucralose and AceK, can modulate NE levels during acute hyperglycemia. When combined with hyperglycemia, both sweeteners counteract the typical reduction in sympathetic activity, leading to increased circulating NE levels. This suggests that non-caloric sweeteners may mitigate the effects of hyperglycemia on sympathetic tone and potentially influence insulin release; it also suggests that, in the presence of glucose, they may activate counterregulatory mechanisms, even though the condition is hyperglycemia rather than hypoglycemia. Overall, these findings underscore the potential of non-caloric sweeteners to alter sympathetic responses during hyperglycemic episodes.

### 4.4. Non-caloric sweeteners (NCS) sucralose and AceK counteract the satiating effect of glucose by increasing food intake after a 12-h fast in rats

Following our investigation into the impact of NCS on glucose sensing and NE levels, we aimed to examine their effects on a parameter regulated by the hypothalamus. Specifically, we focused on refeeding behavior after fasting, as it is a key aspect of the hypothalamic control of feeding responses and energy balance [55].

We first evaluated the effect of NCS on food intake during refeeding in the absence of glucose, comparing these effects to those of the vehicle and glucose alone. The results were presented as the number of pellets consumed after 4 hours following intraperitoneal (i.p.) injection and the reintroduction of food. Rats injected with glucose consumed significantly fewer pellets during the 4-hour refeeding period compared to vehicle-injected rats (Veh. group: 48.63 ± 1.513 mg/g; Glu. group: 38.76 ± 1.148 mg/g, p < 0.001) (**Fig. 4B**). In contrast, injection of sucralose or AceK resulted in significantly higher pellet consumption compared to the vehicle group. Animals in the sucra + veh group consumed more pellets (59.76 ± 1.640 mg/g) than those in the veh group (48.63 ± 1.513 mg/g, p < 0.0001) and those in the glu group (38.76 ± 1.148 mg/g, p < 0.0001). Similarly, animals in the AceK + veh group consumed even more pellets (63.98 ± 1.948 mg/g) than both the vehicle group (p < 0.0001) and the glucose group (p < 0.0001). Food ingestion was comparable between the sucralose and AceK groups (**Fig. 4B**). Over the 4 h period, rats receiving glucose plus sucralose or AceK ate about twice as much as those receiving glucose alone (**Fig. 4C**; p < 0.0001). The increased food intake observed with NCS compared to control groups suggests an orexigenic effect of NCS, while the decreased food intake following glucose administration reinforces the well-established satietogenic effect of glucose in the literature [56].

**Figure 4:**
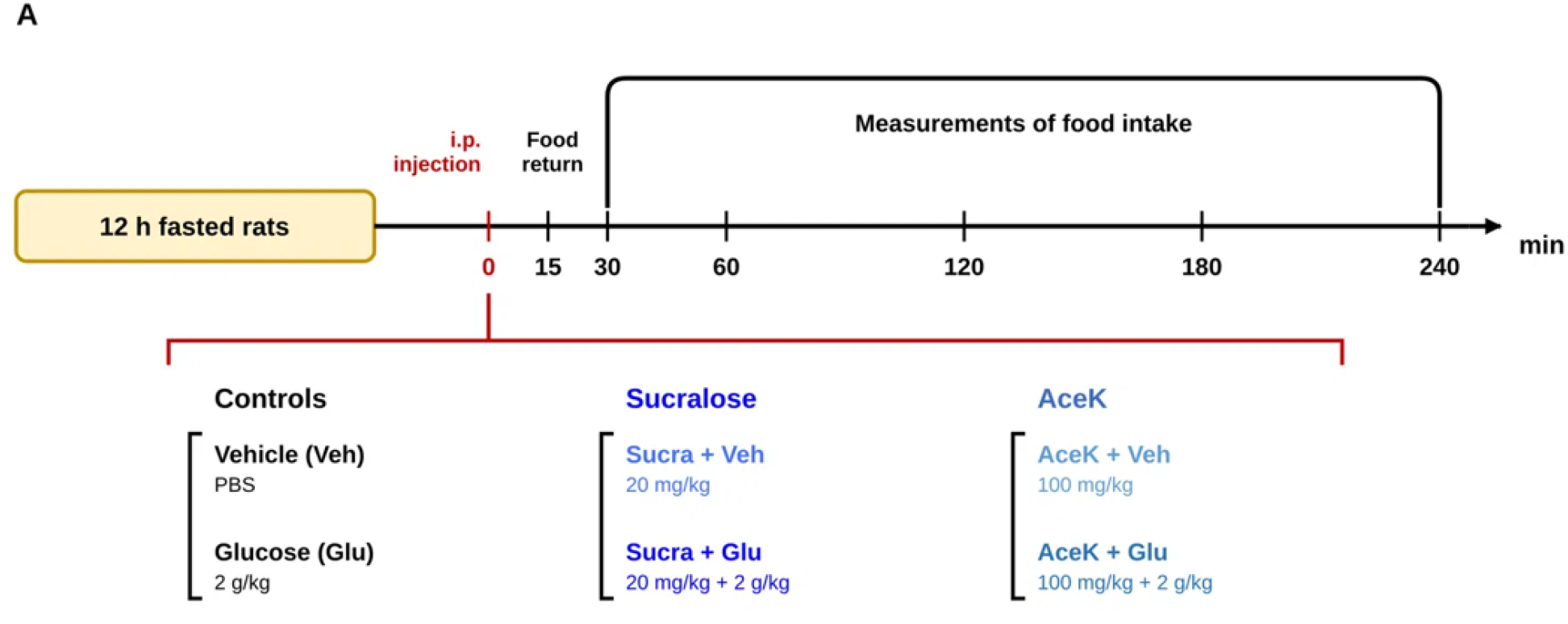
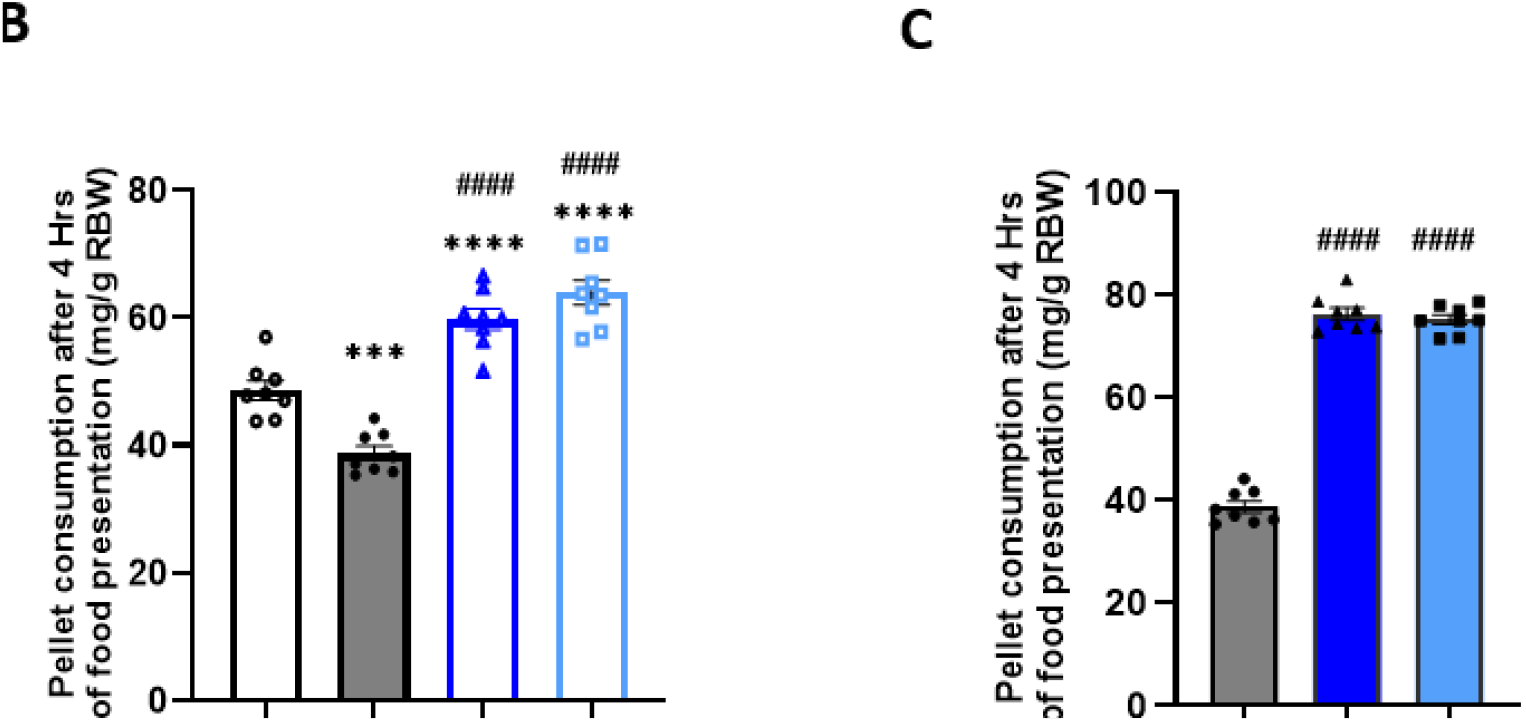
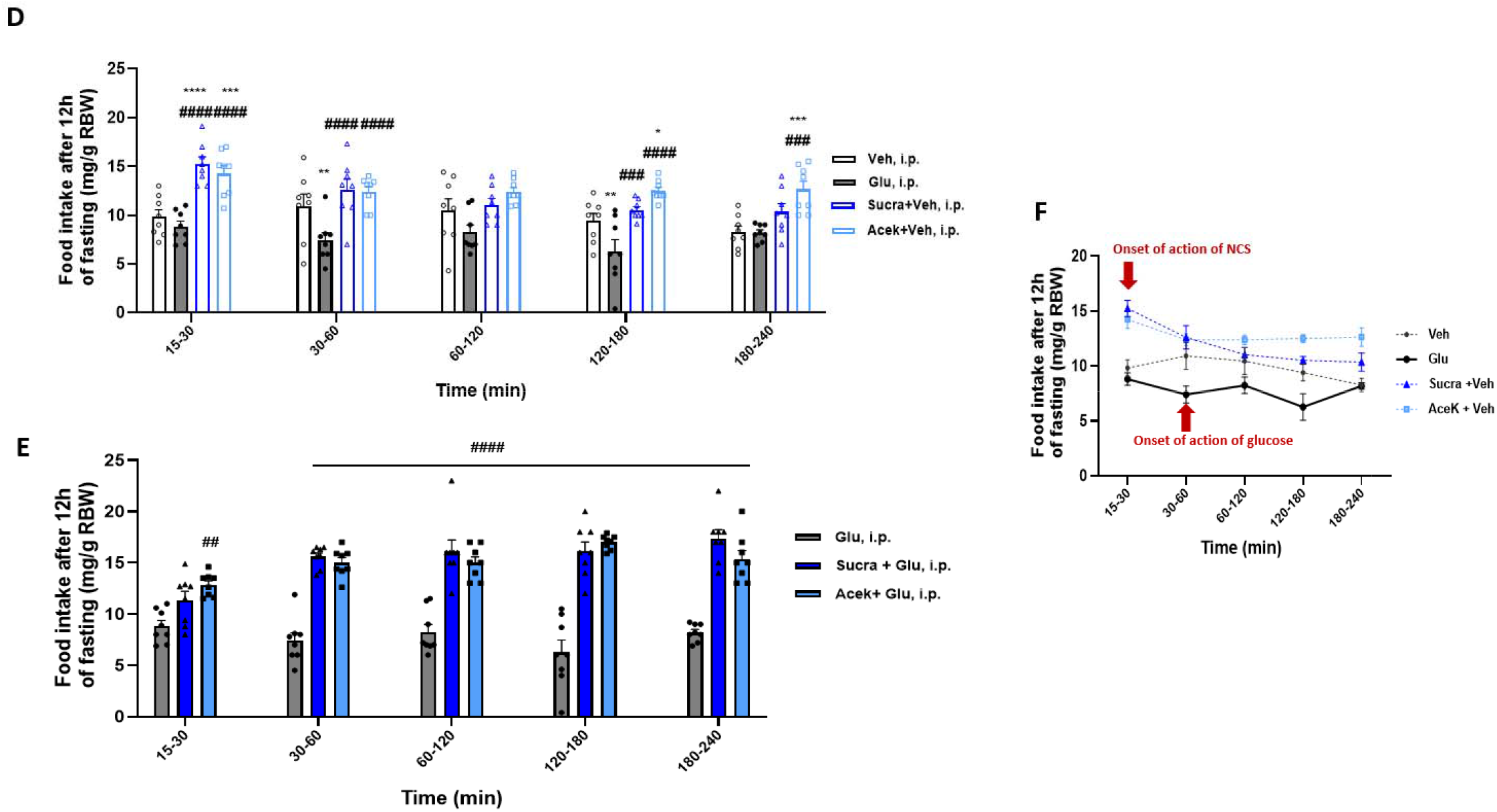
Non-caloric sweeteners increase refeeding after a 12-h fast in rats and counteract the satietogenic effect of glucose. (A) Experimental procedure of the refeeding test after a 12-h fast. Six groups of rats (n = 8 per group) received intraperitoneal (i.p.) injections at time 0. Control groups received vehicle alone (PBS, “Veh”) or glucose (2 g/kg), the positive control for the satietogenic effect of glucose. To assess sucralose, animals received sucralose alone (Sucra + Veh, 20 mg/kg) or glucose (2 g/kg) combined with sucralose (20 mg/kg). To assess AceK, animals received AceK alone (AceK + Veh, 100 mg/kg) or glucose (2 g/kg) combined with AceK (100 mg/kg). Pellets were returned to the cage 15 min after injection to allow the injected substances to enter the bloodstream. Food intake was measured at 30, 60, 120, 180 and 240 min after injection (times counted from the injection). (**B–F**) Food intake, expressed relative to rat body weight (RBW, mg pellets/g). (**B**) Pellet consumption over the 4 h after injection, comparing the vehicle group with groups receiving glucose or NCS alone. (**C**) Pellet consumption in groups receiving NCS combined with glucose, compared with glucose alone. (**D, E**) Feeding intervals, showing the timing of the effects of the injected solutions. (**F**) Food intake curves over time, highlighting the onset of action of NCS compared with glucose (red arrow). Values are mean ± SEM. Statistical significance was assessed by one-way ANOVA followed by Bonferroni’s post hoc test: *p < 0.05, ***p < 0.001, ****p < 0.0001 vs Veh; ####p < 0.0001 vs glucose.

To precisely assess the impact of NCS over time, we analyzed refeeding intervals following injections.

**Figs. 4D and 4F** illustrate the food intake response to the injection of sweeteners and glucose. **Fig. 4F** shows that sweeteners alone have an orexigenic effect that becomes significant within the first 30 min after injection (at 30 min: veh.: 9.825 ± 0.731 mg/g vs. sucra + veh: 15.238 ± 0.739 mg/g, p < 0.0001; AceK + veh: 14.238 ± 0.817 mg/g, p < 0.001). Conversely, the opposite, satiating effect of glucose begins to take effect between 30 and 60 min after injection and persists significantly up to 180 min. This delay is likely due to the time required for its metabolism, unlike NCSs (there was no significant difference between glucose and veh group at this time point with veh.: 9.825 ± 0.731 mg/g vs. glucose: 8.813 ± 0.566 mg/g). Finally, during the last intervals (180-240 min), animals receiving sweeteners continued to consume more food, the orexigenic effect of AceK remaining significant, while the effects of sucralose disappeared completely.

Finally, **Fig. 4E** illustrates the refeeding behavior of animals injected with a combination of glucose and sweeteners (sucralose or AceK) compared to those injected with glucose alone. Animals receiving the combined glucose and sweetener injections consume approximately twice as much food as those injected with glucose alone (p < 0.0001). Especially, within the first 30 min, the AceK + glucose group consumed significantly more food than the glucose one (glucose: 8.813 ± 0.566 mg/g vs. AceK + glucose: 12.850 ± 0.397 mg/g, p < 0.01), an early effect that was not observed in the sucralose + glucose group.Overall, our *in vivo* results suggest that sucralose and AceK significantly alter feeding behavior and autonomic responses compared to glucose alone. Not only do glucose and NCS exhibit opposite effects on food intake after fasting, but the onset of action is also notably more rapid with NCS. Sucralose and AceK prompt a swift increase in food consumption. This rapid orexigenic response contrasts with the delayed satiating effect observed with glucose, highlighting the distinct mechanisms through which NCS and glucose influence appetite and feeding behavior. Furthermore, the combined effect of NCS and glucose shows that NCS abolishes the effect of glucose alone, suggesting that these two pathways are not independent and that the receptor-mediated pathway used by NCS counteracts that of glucose metabolism. It was therefore necessary to determine the nature of these interactions.

### 4.5. Non-Caloric sweeteners decrease dose-dependently ATP production in response to high concentration of glucose, an effect mediated by sweet taste receptors in the hypothalamic cell line GT1-7

To better unravel the effects observed *in vivo*, we switched to a cellular model, the GT1-7 hypothalamic cell line. This cell line was selected due to its established role in previous research on hypothalamic glucose sensing. Derived from mouse hypothalamic tumor tissue, GT1-7 cells have been extensively characterized and shown to exhibit key features of hypothalamic neurons, including responsiveness to glucose stimuli [33,34,35]. First, we investigated the presence of sweet taste receptors and *Gnat3* transcripts in this cell line, then compared transcript levels with those in the mouse hypothalamus (**Fig. 5A**). Both *Tas1r2* and *Tas1r3* mRNAs were present and displayed approximately 4-fold fewer transcripts than mouse hypothalamus (**Fig. 5A**). In contrast, *Gnat3* mRNA levels in the cell line were not significantly different from those measured in the tissue (**Fig. 5A**). These results showed that this line has the cellular features needed to study the T1R pathway. We also verified that the highest sweetener concentrations had no effect on cell viability after 30 min of exposure in 2.5 mM glucose (**Supplementary Fig. S1**). Second, we analyzed the impact of increasing glucose levels from 2.5 mM to 20 mM for 30 min on ATP concentration. A large significant increase in ATP levels was measured (2.5 mM glucose: 6.81 ± 0.62 vs. 20 mM glucose: 25.93 ± 1.82 AU × 1000, p = 0.0002) (**Fig. 5B**). The aim was then to determine whether ATP production in response to an increase in glucose involved sweet taste receptors. To this end, we used gurmarin, a specific inhibitor of these receptors [57]. Under these conditions, we observed no significant difference in ATP production with basal glucose concentration (2.5 mM), nor in response to 20 mM glucose rise, with and without gurmarin (2.5 mM glucose: 6.81 ± 0.62 vs. 2.5 mM glucose + gurmarin: 5.60 ± 0.64; 20 mM glucose: 25.93 ± 1.82 vs. 20 mM glucose + gurmarin: 21.89 ± 2.06 AU × 1000) (**Fig. 5C**). These results suggest that with glucose alone, transduction pathways involving T1Rs sweet receptors have no significant impact on ATP production.

**Figure 5:**
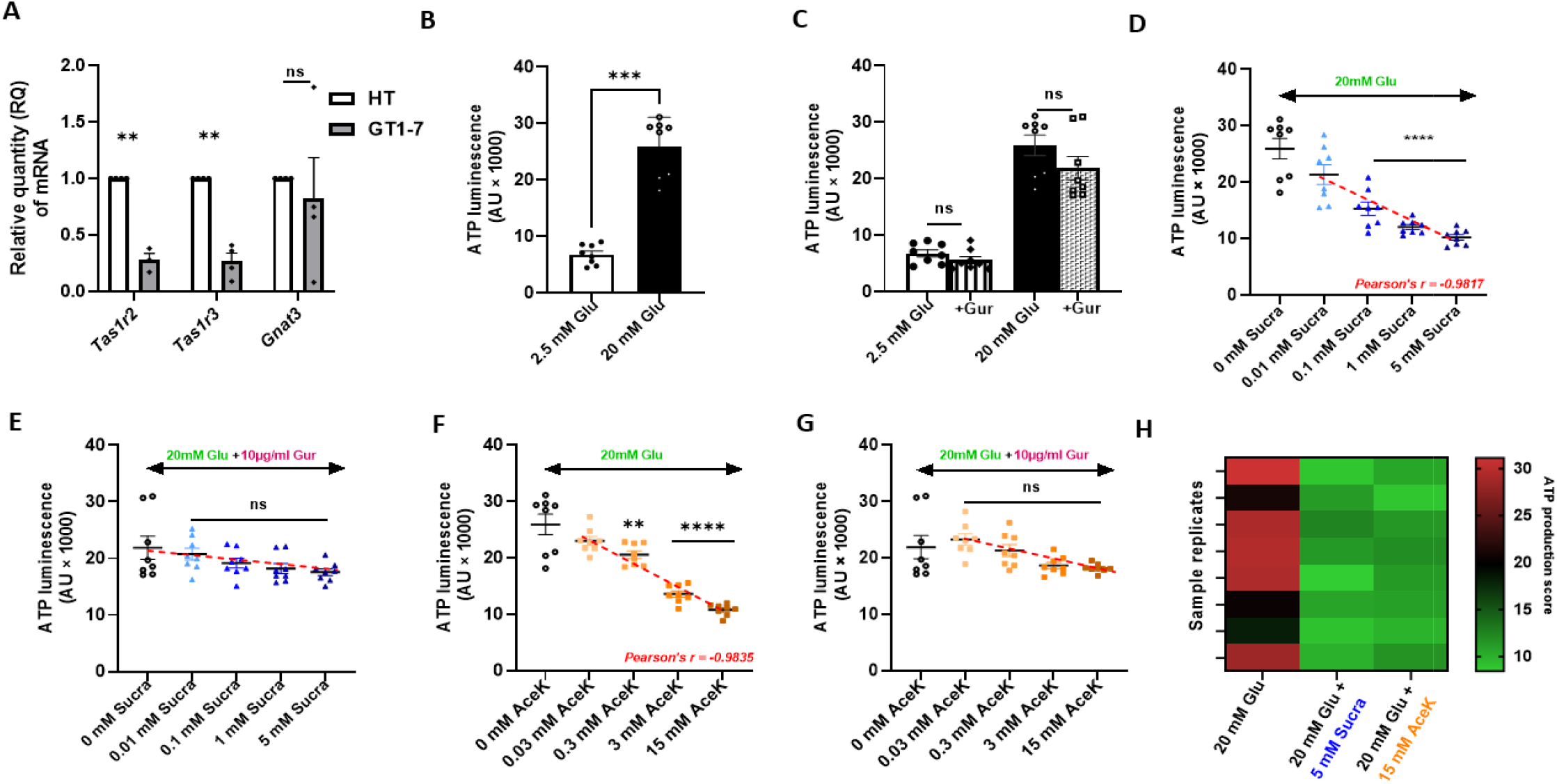
Non-caloric sweeteners induce an acute, dose-dependent decrease in ATP production in GT1-7 hypothalamic cells expressing T1R2/T1R3 and α-gustducin. (**A**) mRNA levels of *Tas1r2*, *Tas1r3* and *Gnat3* measured by RT-qPCR in the GT1-7 cell line and compared with the hypothalamus (HT). Relative quantity (RQ) is the expression of the target genes relative to the reference genes (*Ubc* and *Tbp*). Groups were compared by two-way ANOVA followed by Bonferroni’s post hoc test. Data are mean ± SEM (n = 3–4); *p < 0.05, **p < 0.01. (**B**) ATP concentration after 30 min of incubation with glucose (Glu, 20 mM) vs basal glucose (2.5 mM). (**C**) Effect of gurmarin (Gur), a sweet taste receptor inhibitor, on ATP concentration after 30 min of incubation with 2.5 mM or 20 mM glucose. (**D**) Dose-response effect of increasing sucralose (Sucra) concentrations (0, 0.01, 0.1, 1 and 5 mM) on ATP concentration in cells incubated with 20 mM glucose for 30 min; Pearson’s r is indicated on the graph. (**E**) As in D, in the presence of gurmarin (10 µg/mL). (**F**) Dose-response effect of increasing AceK concentrations (0, 0.03, 0.3, 3 and 15 mM) on ATP concentration in cells incubated with 20 mM glucose for 30 min; Pearson’s r is indicated on the graph. (**G**) As in F, in the presence of gurmarin (10 µg/mL). (**H**) Heat map of ATP concentrations in cells treated with 20 mM glucose alone, 20 mM glucose + 5 mM sucralose, or 20 mM glucose + 15 mM AceK. On the color scale (10 to 30), darker red indicates higher and green lower ATP concentrations. (**B–H**) ATP was measured by luciferin/luciferase luminescence and normalized to the genomic DNA content of each well; values are expressed in arbitrary units (AU) × 1000. Data are mean ± SEM (n = 8 per group; assays in duplicate). Differences were assessed by one-way ANOVA followed by Bonferroni’s post hoc test, or by the Mann–Whitney test. *p < 0.05, **p < 0.01, ***p < 0.001, ****p < 0.0001; ns, not significant

We then proceeded to evaluate the impact of the first Non-Caloric Sweetener (NCS), sucralose, on ATP production in the presence of increased glucose levels (20 mM) for 30 min. Our aim was to mimic *in vivo* conditions of hyperglycemia in the presence of sweetener. Dose-responses were performed in the presence of elevated glucose with increasing concentrations of sucralose (0.01, 0.1, 1 and 5 mM) to quantify ATP production (**Fig. 5D**). Under these conditions, we observed an inverse correlation between increasing concentrations of sucralose and ATP production starting from a concentration of 0.1 mM (**Fig. 5D**). Significant declines in ATP levels were noted from this concentration onwards, compared to the sucralose-free group (p < 0.0001). The correlation was highly significant with **Pearson’s r = −0.9817**, p = 0.003. The question arose whether the reduction in ATP production could be attributed to the binding of sucralose to sweet taste receptors. To address this, we reintroduced gurmarin, a highly specific sweet taste receptor inhibitor. Our results showed that, with gurmarin present, there were no more significant differences in ATP production across the various sucralose concentrations used (**Fig. 5E**).

Subsequently, we conducted analogous experiments using the alternative sweetener acesulfame potassium (AceK), which has been used *in vivo*. Given that AceK is known for its lower sweetness potency—approximately one-third that of sucralose—we employed a concentration three times higher to match the sweetening power of sucralose. The results with AceK exhibited a similar trend to those observed with sucralose, showing a comparable inhibitory pattern in ATP production (**Fig. 5F**, 5G). The Pearson correlation coefficient for this relationship was **r = −0.9835**, with a significance level of p = 0.0025. To summarize the major findings of this experiment, **Fig. 5H** presents a quantitative, color-coded visualization of ATP production measured during a 30-min glucose incubation. In summary, gurmarin did not alter ATP production with glucose alone, but only the decrease induced by sucralose and AceK, which was no longer significant in its presence, demonstrating that T1R2/ T1R3 are involved.

### 4.6. Non-Caloric sweeteners enhance ROS generation in GT1-7 hypothalamic cell line under hyperglycemic conditions

The results indicating a decrease in ATP production suggested that ROS generation might also be impacted. In normal physiology, as shown in **Fig. 6A**, hyperglycemia led to an increase in ROS generation. When comparing the effects of different glucose concentrations, we observed that after 10 min, ROS levels were significantly higher with 20 mM glucose (16,866 ± 3,594 AU) than with 2.5 mM glucose (2,740 ± 1,377 AU, p < 0.05). The difference was even more pronounced at 15 min (20 mM: 29,489 ± 6,958 AU vs. 2.5 mM: 11,267 ± 2,595 AU, p < 0.01), indicating that higher glucose levels substantially elevated ROS production over time.

**Figure 6:**
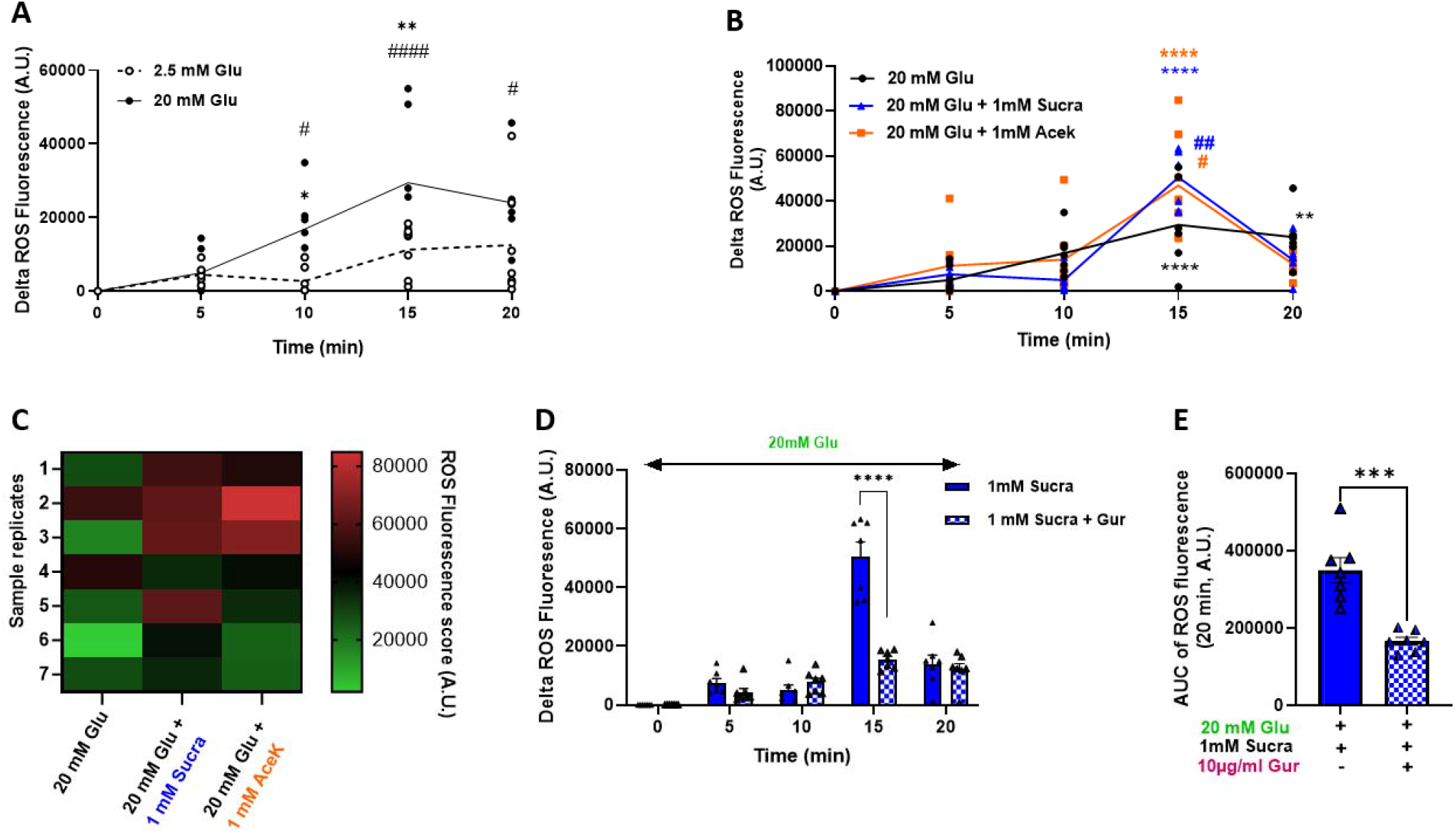
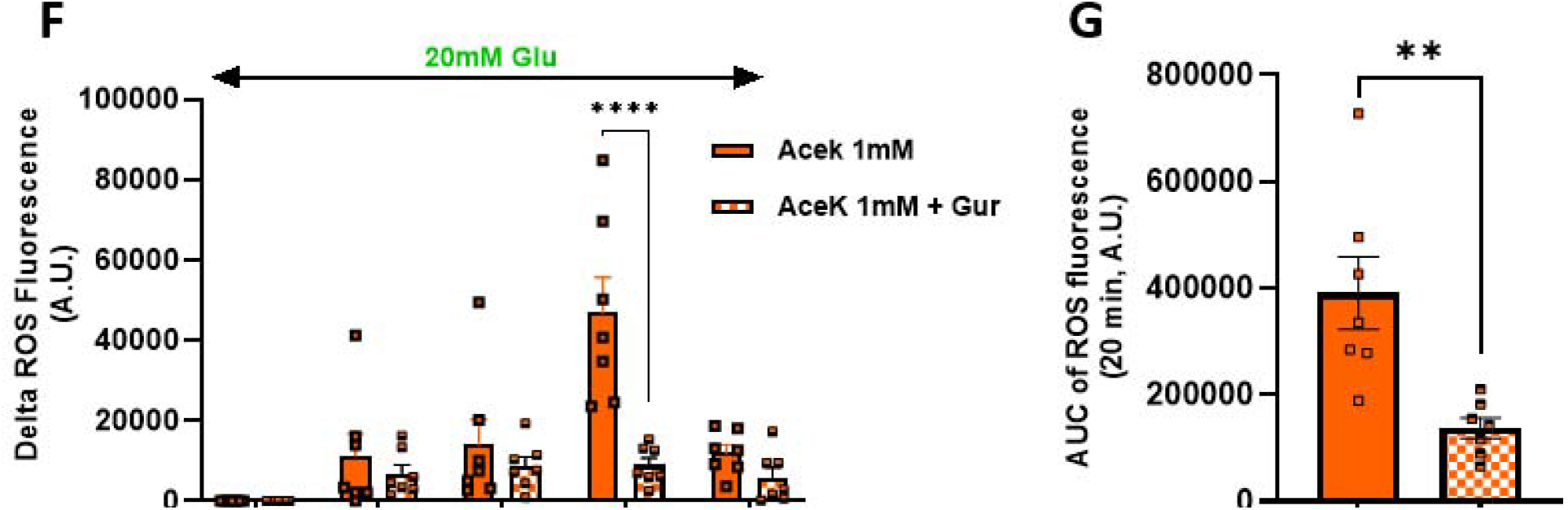
Non-caloric sweeteners induce acute increases in ROS generation in GT1-7 hypothalamic cells expressing T1R2/T1R3 and α-gustducin. ROS generation was measured as extracellular H₂O₂ release (Amplex Red). (**A**) ROS production over 20 min with 2.5 mM vs 20 mM glucose. (**B**) ROS generation curves after the addition of 1 mM sucralose or AceK. (**C**) Heat map summarizing the ROS generation results across conditions. (**D**) Effect of gurmarin on sucralose-induced ROS generation in the presence of glucose over 20 min. (**E**) Area under the curve (AUC) of ROS generation in the sucralose groups with and without gurmarin. (**F**) ROS generation with AceK in the presence of glucose over 20 min. (**G**) AUC of ROS generation in the AceK groups. Statistical analyses were performed with two-way ANOVA and the Mann–Whitney test (n = 7 per group). Values are the change from baseline at the indicated time, expressed as mean ± SEM. *, comparison within the same group between different times; #, comparison with the control and glucose groups.

Next, we examined the influence of NCS on ROS generation, as shown in **Fig. 6B**. At t = 15 min, the impact on ROS generation became evident, with a rise in both groups treated with NCS. Notably, sucralose had a more pronounced effect on ROS generation compared to AceK. Specifically, at t = 15 min, ROS generation in the 20 mM glucose group was 29,489 ± 6,958 AU, compared to 50,530 ± 4,946 AU in the sucralose with 20 mM glucose group (p < 0.01) and 46,880 ± 8,721 AU in the AceK with 20 mM glucose group (p < 0.05). This effect was observed only at t = 15 min and returned to the same levels as hyperglycemia starting from t = 20 min. These results were well visualized in the heat map of **Fig. 6C**, which showed a greener color in the glucose-only group compared to the red in the NCS-treated groups, indicating higher ROS generation with NCS.

To further explore the role of sweet taste receptors in the observed increase in ROS generation, we evaluated the effect of gurmarin, a known antagonist of sweet taste receptors. By adding gurmarin to the same conditions, we aimed to determine if the ROS generation induced by NCS was mediated through these receptors. As shown in **Fig. 6D**, the impact of sucralose on ROS generation at 15 min was significantly reduced in the presence of gurmarin. Specifically, ROS levels at 15 min were markedly lower in the sucralose + gurmarin group (15,384 ± 1,152 AU) compared to the sucralose-only group (50,530 ± 4,946 AU), with a p-value < 0.0001. Additionally, the area under the curve (AUC) for ROS generation over 20 min, as depicted in **Fig. 6E**, confirmed that the presence of gurmarin significantly diminished the ROS generation associated with sucralose, indicating that the observed effect was indeed mediated through sweet taste receptors.

Similarly, we observed a consistent trend with AceK (**Fig. 6F**). The addition of gurmarin to the AceK treatment conditions led to a significant reduction in ROS generation. At 15 min, the ROS in the AceK + gurmarin group (8,947 ± 1,789 AU) were markedly lower compared to those in the AceK only group (46,880 ± 8,721 AU), with a p-value < 0.0001. This trend was further confirmed by the area under the curve (AUC) analysis over 20 min, as shown in **Fig. 6G**, which illustrated that the presence of gurmarin effectively mitigated the ROS generation induced by AceK.

Our findings demonstrated that non-caloric sweeteners (NCS), specifically sucralose and acesulfame potassium (AceK), significantly elevated reactive oxygen species (ROS) generation in the GT1-7 hypothalamic cell line. This increase in ROS production was notably influenced by the presence of glucose, indicating a synergistic effect when NCS were combined with hyperglycemic conditions and this effect was mediated by sweet taste receptors.

### 4.7. Non-nutritive sweeteners sucralose and AceK reduce stimulated substrates-driven mitochondrial respiration in the hypothalamic GT1-7 cell line

The observed decrease in ATP production in the presence of a high glucose plus sweetener combination could be due to its impact on mitochondrial respiration, as well as the increase in reactive oxygen species (ROS) generation. To investigate these possibilities, we conducted experiments to assess mitochondrial oxygen consumption under both basal and stimulated respiration conditions. These conditions were designed to mimic those used for measuring ATP production, thereby allowing us to evaluate how the combination of high glucose and non-caloric sweeteners affects mitochondrial function and ROS production.

We first assessed the impact of increasing sucralose concentrations (0.001, 0.01 and 1 mM) on basal respiration in GT1-7 cells. Under these conditions, we measured no significant effect of sucralose at any concentration on cell oxygen consumption (**Fig. 7A**). In contrast, when respiration was previously stimulated by substrate-driven respiration (SDR) – mimicking reduced equivalents induced by glucose in the complexes – sucralose (1 mM) significantly decreased mitochondrial respiration in GT1-7 cells (21.80 ± 1.18 vs. 17.08 ± 1.18 pmol O_2_/(s*M.cells), p< 0.05) (**Fig. 7B**).

**Figure 7:**
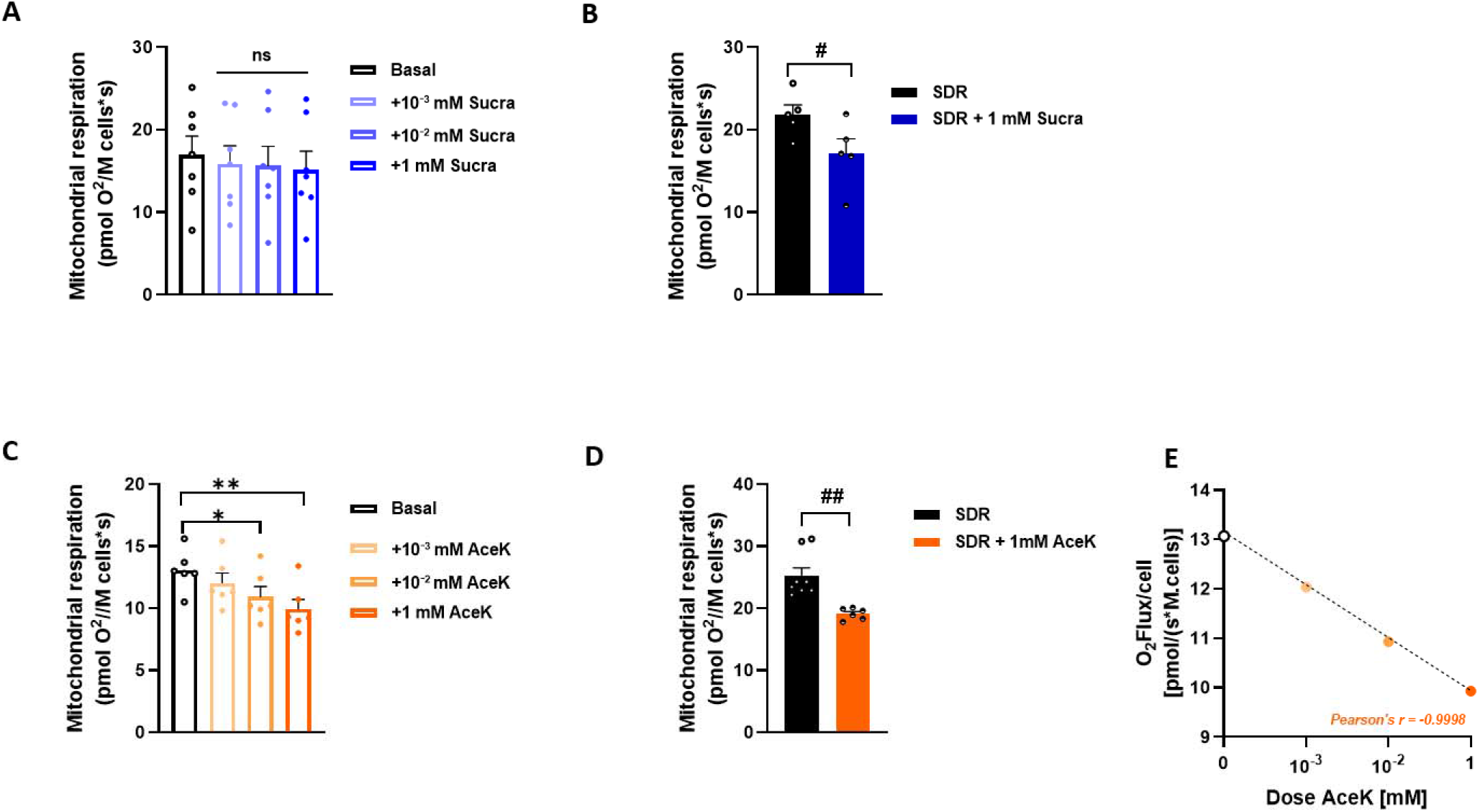
Effects of non-caloric sweeteners on mitochondrial respiration in GT1-7 hypothalamic cells. (**A–D**) Oxygen consumption of saponin-permeabilized GT1-7 cells measured by high-resolution respirometry. Basal respiration was measured in the absence of exogenous substrates; substrate-driven respiration (SDR) corresponds to the maximal OXPHOS stimulation after addition of succinate and ADP. (**A**) Effect of increasing sucralose (Sucra) concentrations (10^−3^, 10^−2^ and 1 mM) on basal respiration. (**B**) Effect of sucralose (1 mM) on SDR. (**C**) Effect of increasing AceK concentrations (10⁻³, 10⁻² and 1 mM) on basal respiration. (**D**) Effect of AceK (1 mM) on SDR. (**E**) Relationship between AceK concentration and basal oxygen consumption (Pearson correlation). Oxygen flux is expressed in pmol O₂/ (s × 10^6^ cells). Data are mean ± SEM (n = 7 per condition). Statistical analysis was performed with a mixed-effects model (REML; treatment as fixed effect, individual preparation as random effect) followed by Bonferroni’s post hoc test, and a paired t-test for B and D. *p < 0.05 and **p < 0.01 vs basal; #p < 0.05 and ##p < 0.01 vs SDR; ns, not significant.

These experiments were repeated with AceK. Here, increasing concentrations of the sweetener AceK (0.001, 0.01 and 1 mM) significantly decreased basal mitochondrial respiration from 10^−2^ mM concentration (10^−2^ mM AceK: 10.93 ± 0.81 vs. basal state: 13.06± 0.67 pmol O /(s*M.cells), p< 0.05 and 1 mM AceK: 9.93 ± 0.77 vs. basal state: 13.06± 0.67 Acepmol O_2_/(s*M.cells), p<0.01) (**Fig. 7C**). This decrease in basal O_2_ consumption was dose-dependent, as depicted in **Fig. 7E** and confirmed by Pearson correlation analysis, with a negative correlation between AceK concentration and oxygen consumption (r= −0.9998, p<0.0002). Under conditions of stimulated mitochondrial respiration (SDR), the inhibitory effect of AceK (1 mM) on O_2_ consumption was still significantly displayed (SDR: 25.24± 1.27 pmol O_2_/(s*M.cells) vs. SDR + 1 mM AceK: 19.1 ± 0.37 pmol O_2_/(s*M.cells), p<0.01) (**Fig. 7D**). Altogether, these data show that the two sweeteners sucralose and AceK significantly reduce oxidative phosphorylation under stimulated energetic conditions (and even under basal conditions for AceK), which could explain the drop in ATP production measured in the GT1-7 hypothalamic cell line.

## 5 | DISCUSSION

This study provides new evidence that non-caloric sweeteners (NCS), specifically sucralose and acesulfame potassium (AceK), can acutely alter central glucose signaling in hypothalamic neurons involved in energy balance. When delivered toward the brain together with a stimulating dose of glucose, these compounds, commonly consumed under the assumption of metabolic neutrality, blunted hypothalamic glucose-induced insulin secretion and reversed the glucose-induced fall in plasma norepinephrine. Given intraperitoneally, they increased food intake during a refeeding test and counteracted the satiating effect of glucose. In hypothalamic GT1-7 cells, sucralose and AceK reduced glucose-induced ATP production and increased hydrogen peroxide generation in a manner prevented by the selective sweet taste receptor inhibitor gurmarin, and they reduced mitochondrial respiration. By delivering the compounds through the carotid artery (brain, to leave peripheral blood glucose level unchanged) or intraperitoneally, which bypasses systemic circulation or the digestive tract respectively, our study addresses an important mechanistic gap in understanding how NCS may influence metabolic regulation. Altogether, these findings suggest that the simultaneous presence of an NCS and a rise in glucose levels (as might occur during a meal accompanied by a sweetened beverage) disrupts hypothalamic glucose sensing and oxidative metabolism through sweet taste receptors, which may contribute to a defect in the physiological control of metabolism.

Our gene expression mapping showed that the sweet taste receptor subunits and signaling components are expressed in the rat hypothalamus, with distinct regional profiles. Among brain regions, *Tas1r2* and *Gnat3* mRNAs were most abundant in the hypothalamus, whereas *Tas1r3* and *Trpm5* were highest in the cerebellum (*Tas1r3* at about 250% of hypothalamic levels). Hypothalamic levels were lower than in peripheral taste-related tissues: about three-fold lower than in the tongue for *Tas1r2*, six-fold for *Tas1r3*, five-to seven-fold for *Trpm5* and more than 35-fold for Gnat3. This pattern extends earlier reports in mice [58; 27] and is in line with human RNA-seq data (**Supplementary Fig. S2**), supporting the hypothalamus as a central site of sweet taste receptor expression and raising the possibility of broader roles for T1R3 in the brain. Recent reviews similarly describe the sweet taste receptor as a potential regulator of metabolism in extra-oral tissues, including the brain [48;59;60]. Sweet taste signaling may therefore have been repurposed as a central energy-sensing input layered on top of the canonical metabolic glucose-sensing pathway and, as we show below, capable of interfering with it.

In physiological terms, acute intracarotid injection of a stimulatory dose of glucose toward the brain triggers a rapid insulin secretory response, peaking at 1 min, via hypothalamic glucoregulatory circuits, activating glucose-excited (GE) neurons, as previously shown in both rats and mice [20; 21; 22; 23]. When sucralose (1 mM) or AceK (3 mM) was co-injected with this stimulating glucose dose, the insulin peak was significantly blunted, by about a quarter with sucralose and more than a third with AceK, whereas neither sweetener, injected with basal glucose, triggered an insulin peak. Peripheral glycemia remained unchanged in every group. The injected concentrations are of the order of the EC_50_ values of sucralose and AceK at the sweet taste receptor T1R2/T1R3 (about 0.23 and 1.3 mM in HEK293T cells; [61]), before dilution in blood. These values are roughly 130-to 430-fold (sucralose) and 20-to 75-fold (AceK) lower than the 30 to 100 mM range at which sugars activate the receptor [62; 63; 64], although these ratios combine values from different assays and should be read as orders of magnitude. The most parsimonious explanation is therefore that these NCS interfere with the canonical glucose-sensing pathways of glucose-responsive hypothalamic neurons, distorting the downstream signal that normally reflects glucose availability.

To our knowledge, this is the first demonstration of a direct, acute central effect of this kind, consistent with a receptor-mediated mechanism. It addresses an ambiguity left open by earlier human work: Pepino et al. [65] reported that sucralose ingestion alters glycemic and hormonal responses to an oral glucose load, and Gómez-Arauz et al. [66] later showed that a single 48 mg sucralose sip is enough to raise serum insulin during an oral glucose tolerance test (OGTT) in healthy adults, but they acknowledged, as a limitation of their study, a possible role of the sweet taste of sucralose in modulating insulin secretion via the central nervous system. Our intracarotid design was built precisely to isolate that central component, and our data suggest that it is sufficient to blunt glucose-induced insulin secretion.

Beyond insulin secretion, NCS significantly modulated autonomic outflow during acute central hyperglycemia (elevation of brain glucose) induced by intracarotid glucose injection. Elevated brain glucose alone lowered plasma norepinephrine (NE) 10 min after injection, consistent with reduced sympathetic activity [67] and with a shift toward parasympathetic (vagal) dominance, which favours insulin secretion [68]. When sucralose or AceK was co-injected with the stimulating glucose dose, this decrease was abolished and plasma NE instead rose, whereas the sweeteners given with basal glucose did not change NE. This suggests that the sweeteners do not directly activate sympathetic outflow but rather disrupt glucose-dependent autonomic regulation. To our knowledge, this has not been reported previously, and it is in line with the established role of hypothalamic glucose sensing in controlling parasympathetic and sympathetic outflow to the endocrine pancreas [69;23]; our data suggest that NCS can disrupt this central regulation. The combination of blunted, putatively vagally mediated insulin secretion and increased sympathetic activity offers a plausible acute basis for the metabolic dysregulation reported after chronic NCS exposure in both animal and human studies.

Behaviorally, 12-hour-fasted rats injected intraperitoneally with sucralose or AceK ate more over 4 hours than vehicle-injected rats, whereas glucose reduced re-intake, consistent with its well-established satiating effect [42; 22]. When combined with glucose, the sweeteners prevented this satiating effect and rats receiving glucose plus a sweetener ate roughly twice as much as those receiving glucose alone, with a faster onset for AceK. This orexigenic increased response with sweeteners occurred within 30 min, preceding the satiety phase induced by glucose alone, which became apparent between 30 and 60 min and persisted up to 180 min (**Figs. 4D, 4F**). Overall intake was similar with the two sweeteners, but the effect of AceK persisted until the last interval (180 to 240 min) whereas that of sucralose had disappeared.

These observations are compatible with the sweet-calorie uncoupling hypothesis of Swithers and Davidson (2008) [70], whereby sweet taste without caloric content weakens the learned association between sweetness and energy intake and promotes compensatory eating. However, that framework relies on repeated exposure and learning, whereas the effect we observed was acute, which points to an immediate mechanism. Earlier work emphasized the chronic effects of NCS on appetite, body weight and glucose tolerance [71;72]; here our data show that a single acute exposure is already sufficient to measurably shift feeding behavior, underlining the sensitivity of appetite-regulating circuits to sweet, energy-free signals.

To probe the mechanisms underlying these *in vivo* effects, we turned to the mouse hypothalamic GT1-7 cell line, which expresses glucose-sensing machinery [32;73;74]. In these cells, *Tas1r2* and *Tas1r3* mRNAs were present at about a quarter of mouse hypothalamic levels, and *Gnat3* at similar levels. Raising glucose from 2.5 to 20 mM increased ATP about four-fold, and gurmarin did not significantly alter this rise. Both sucralose (from 0.1 mM) and AceK (from 0.3 mM) dose-dependently reduced the glucose-induced ATP increase, an effect that was no longer significant in the presence of gurmarin, supporting a role for T1R2/T1R3 signaling in the metabolic disruption.

This apparent paradox, a non-caloric ligand overriding the effect of an abundant, caloric one, could be more readily explained if glucose and NCS act on largely distinct components of the signaling network, and with very different potencies at T1R2/T1R3, rather than competing for a unique similar pathway. Our data show that gurmarin, an inhibitor of rodent T1R2/T1R3 sweet taste receptors, did not significantly affect the glucose-evoked increase in intracellular ATP. This indicates that glucose-induced ATP production runs chiefly through the canonical, receptor-independent route (uptake via glucose transporters, followed by glycolysis and mitochondrial oxidative phosphorylation, K_ATP_ channels, ROS, etc.) and that T1R2/T1R3 signaling is not required for it. This contrasts with Kohno et al. [27], who showed that gurmarin suppressed glucose-evoked calcium responses in arcuate neurons, but the readouts differ (calcium activity versus ATP production).

By contrast, sucralose and AceK are not metabolic substrates, yet they are far more potent agonists of the sweet taste receptor T1R2/T1R3 than glucose [75;62]. Their EC_50_ values in heterologous cells are about 0.23 and 1.3 mM, respectively, against tens to hundreds of millimolar for sugars [61;63], in line with their sweetness potencies of about 600– and 200-fold that of sucrose [61], and low-molecular-weight sweeteners bind within the ligand-binding cleft of the T1R2 extracellular domain [41]. Sucralose and AceK reduced glucose-induced ATP in our GT1-7 cells from 0.1 and 0.3 mM, respectively. T1R2/T1R3 signals through Gα-gustducin and PLCβ2-mediated calcium release, the canonical sweet taste transduction cascade [63], whose components are expressed in hypothalamic neurons, including the arcuate nucleus (ARC) [58; 27]. Sub-millimolar to low-millimolar concentrations of a high-potency NCS may therefore be sufficient to strongly engage this cascade, whereas glucose recruits it only weakly, mainly at high concentrations [27;28]. A plausible mechanism is that the resulting calcium/second-messenger signal perturbs mitochondrial function, possibly through altered mitochondrial calcium handling, membrane potential or electron transport chain activity, thereby lowering ATP production and increasing reactive oxygen species (ROS) despite normal glucose delivery and glycolytic metabolism [76;77]. Consistent with a sweet taste receptor-dependent ROS response to NCS, sucralose increases ROS through T1R3 in hepatic cells [78], increases ROS in mesenchymal stromal cells [79], and NCS induce ER stress in hypothalamic cells [45]. Under this hypothesis, when glucose levels alone rise, the contribution of T1R2/T1R3 signaling is negligible and does not interfere with the mitochondrial metabolic pathway. We hypothesize that T1R2/T1R3 signaling is only recruited under marked hyperglycemia, as a protective mechanism against excessive glucose oxidation. In the presence of a high-potency NCS, however, this pathway would be strongly activated even at non-critical glucose levels, possibly counteracting glucose oxidation by inhibiting mitochondrial respiration and increasing ROS, as suggested by our mitochondrial respiration results.

The role of ROS in hypothalamic glucose sensing is double-edged: at low levels, ROS act as signaling molecules, whereas excess ROS impairs mitochondrial function and neuronal excitability [20;76;80]. Here, both sucralose and AceK (1 mM) transiently increased hydrogen peroxide generation in GT1-7 cells exposed to 20 mM glucose, with a peak at 15 min, and this increase was prevented by gurmarin, supporting the involvement of T1R2/T1R3 in redox disruption. This rise in ROS was accompanied by a drop in ATP and by reduced mitochondrial respiration. In permeabilized GT1-7 cells, both sweeteners reduced substrates-stimulated oxygen consumption, and AceK also lowered basal respiration in a dose-dependent manner from 0.01 mM. These data suggest that T1R2/T1R3 does not contribute to glucose-driven ATP production but may instead modulate how much glucose is oxidized once inside the cell, acting as a negative-feedback sensor. This uncoupling between glucose perception and glucose oxidation creates a functional vulnerability that a high-potency, non-caloric sweet ligand such as an NCS can exploit.

Overall, our data support a model in which NCS-induced activation of central sweet taste receptors uncouples glucose sensing from energy metabolism (**Fig. 8**). Glucose is available and would normally support ATP synthesis and neuronal activation, yet the concomitant presence of an NCS distorts this signal, producing a paradoxical cellular state (reduced mitochondrial respiration with low ATP and high oxidative stress) that is incompatible with normal glucose sensing and, more generally, with energy supply to excitable neurons. A large proportion of glucose-excitable neurons have been described as becoming inhibited when glucose concentration drops, due to decreased ATP production and the resulting opening of the K_ATP_ channel (triggering a signal classically associated with hypoglycemia). However, blood glucose levels rise here. Yet, because ATP production decreases in the presence of noncaloric sweeteners, the physiological responses are those typically seen in response to hypoglycemia: increased norepinephrine release, decreased insulin secretion, and increased food intake. While earlier studies focused largely on peripheral tissues and chronic exposure models, our work shows that acute exposure to an NCS, in the presence of glucose, is sufficient to induce rapid autonomic, metabolic and behavioral disorders.

**Figure 8:**
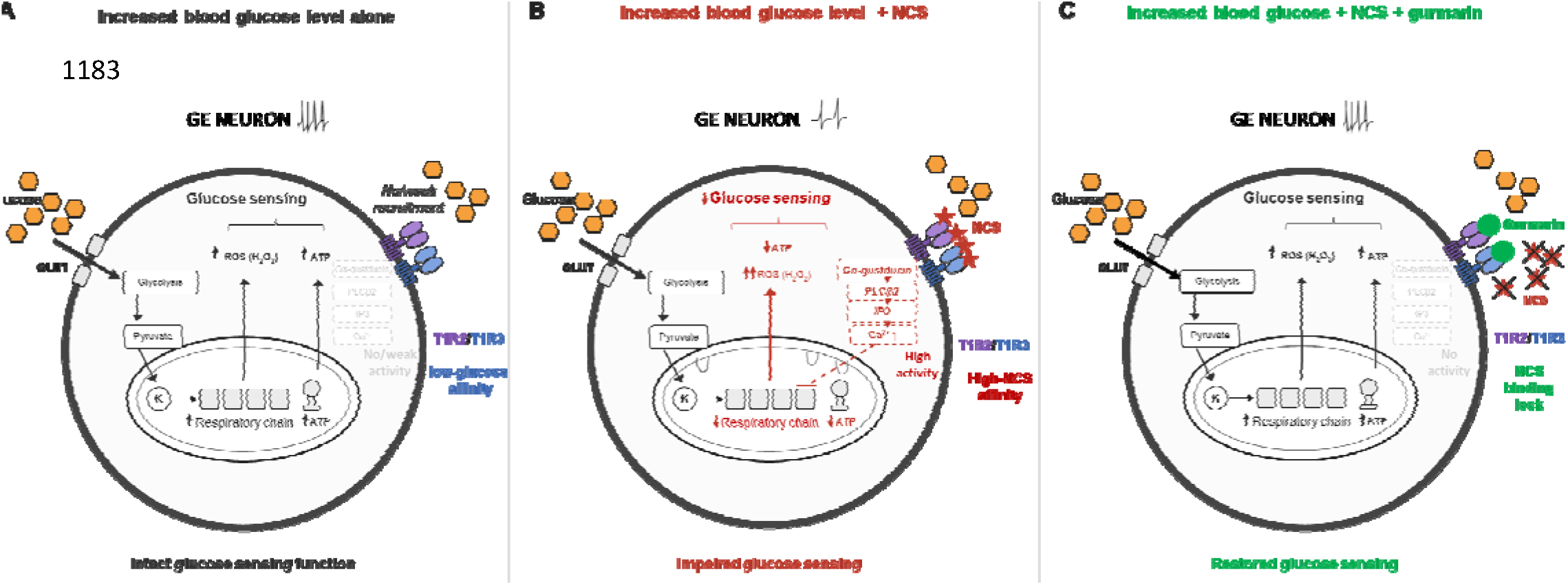
Proposed model of NCS-induced disruption of hypothalamic glucose sensing in gluco-excited (GE) neurons. (**A**) High glucose alone: glucose enters through GLUT transporters and is converted by glycolysis to pyruvate, which enters the mitochondria to fuel the Krebs cycle and the respiratory chain, raising ATP and ROS (H₂O₂). These would support neuronal depolarization and glucose sensing through, at least, KATP and TRPC3 channels. T1R2/T1R3 is weakly recruited by glucose. (**B**) High glucose + NCS: NCS, which are far more potent than glucose at T1R2/T1R3, are proposed to trigger Gα-gustducin/PLCβ2/IP₃-dependent calcium signaling (dashed) that decreases mitochondrial respiration. ATP is reduced despite glucose availability and H₂O₂ is abnormally elevated, which would impair glucose sensing. (**C**) High glucose + NCS + gurmarin: blocking T1R2/T1R3 with gurmarin reduces the NCS-induced changes in ATP and H₂O₂, which would preserve glucose sensing. Solid elements, effects observed in this study; dashed elements, proposed mechanism (neuronal depolarization was not measured).

Several limitations should be considered when interpreting these results. First, all *in vivo* experiments were performed in male rats; given the known sex differences in hypothalamic glucose sensing and in sweet taste sensitivity, extending this work to females will be a priority. Second, the doses and routes used here (intracarotid and intraperitoneal) were chosen to isolate a central mechanism rather than to mimic habitual dietary exposure; the intraperitoneal injections of the refeeding test are systemic, and the dose equivalence with everyday NCS consumption remains to be established. Third, GT1-7 cells are an immortalized mouse hypothalamic cell line that expresses *Tas1r2* and *Tas1r3* at about a quarter of hypothalamic levels and cannot fully reproduce the heterogeneity of native ARC neurons.

These acute mechanistic findings are also consistent with recent epidemiological and policy evidence. In an 8-year prospective cohort, higher habitual consumption of low– and no-calorie sweeteners was associated with faster cognitive decline, including for aspartame, saccharin and acesulfame-K, although not in participants aged 60 years and over [81]; sucralose was not among the sweeteners assessed. In 2023, the World Health Organization issued a conditional recommendation against the use of non-sugar sweeteners to control body weight or reduce the risk of noncommunicable diseases, based on the absence of a demonstrated long-term benefit and on signals of possible undesirable effects [82]. Our data do not address cognition or long-term outcomes, and these associations are observational and may be confounded. Nevertheless, they provide a concrete, receptor-dependent neuronal mechanism that could plausibly act upstream of such chronic associations.

Collectively, these results suggest that sweet-taste receptors in the brain help modulate the response of glucose-excited neurons by dose-dependently reducing their signaling when glucose levels are too high, which could represent a protective mechanism. By activating these receptors as blood glucose levels rise normally, NCSs generate a misleading signal that disrupts mitochondrial function and reduces energy (ATP) production, which is essential for an adequate response. Sweeteners are designed to provide an intense sweet taste, which relies on an affinity for the T1R2/T1R3 receptors that is much higher than that of glucose. This suggests that NCSs with an affinity similar to that of glucose should be used to restrict these disruptions. This new hypothesis needs to be tested, as it could represent a promising avenue for limiting the undesirable metabolic effects of current sweeteners.

## 6 | CONCLUSION

This study shows that the non-caloric sweeteners, sucralose and acesulfame potassium can acutely disrupt central glucose sensing through sweet taste receptors. We first showed that the sweet taste receptor subunits and signaling components are expressed in the rat hypothalamus. *In vivo*, NCS delivered toward the brain with a stimulating dose of glucose blunted glucose-induced insulin secretion and abolished the associated fall in plasma norepinephrine. Given systemically, they increased food intake and counteracted the satiating effect of glucose. In hypothalamic GT1-7 cells, NCS reduced glucose-induced ATP production, increased reactive oxygen species and decreased mitochondrial respiration, with the ATP and ROS effects prevented by the sweet taste receptor inhibitor gurmarin. Together, these results support a model in which high-potency activation of central sweet taste receptors by NCS uncouples glucose sensing from energy metabolism, generating a misleading signal despite normal glucose availability. This provides a receptor-dependent mechanism that may contribute to the metabolic disturbances associated with NCS consumption, and it suggests that sweeteners with a potency closer to that of glucose could limit these effects.

## Supporting information

Supplementary Table S1 TaqMan assays

Supplementary appendix Gurmarin

## 7 | ACKNOWLEDGMENTS

We extend our deepest gratitude to the staff at our animal care facilities for their meticulous attention and care. Their dedication to maintaining the well-being of the animals was crucial to the success of this study, and we greatly appreciate their invaluable support and expertise.

## 10 | AUTHOR CONTRIBUTIONS

According to the CRediT Taxonomy guidelines: **J.H**. contributed to formal analysis (lead), investigation (lead), methodology (equal), validation (lead), visualization (equal), and writing original draft (equal). **C.F**. assisted with formal analysis (equal), investigation (equal), methodology (equal), validation (equal), visualization (equal), and supervision (equal). **B.A**. assisted with formal analysis (equal), investigation (equal), methodology (equal), **F.L**. conducted investigation (equal), resources (equal), and validation (equal). **S.G**. performed investigation (equal), resources (lead), validation (equal), **L. B**. and **G.W**. contributed to reviewing and editing (equal). **C.L**. assisted with conceptualization (lead), formal analysis (lead), funding acquisition (lead), investigation (equal), methodology (equal), project administration (lead), supervision (lead), validation (lead), visualization (lead), writing—original draft (lead), reviewing and editing (lead).

