## Supplementary figures and images for "Non-caloric sweeteners combined with glucose affect hypothalamic glucose sensing-induced insulin secretion and refeeding through neuronal cellular metabolism: *in vivo* and *in vitro* approaches"

### Supplemental Figure S1

#### Slide 1
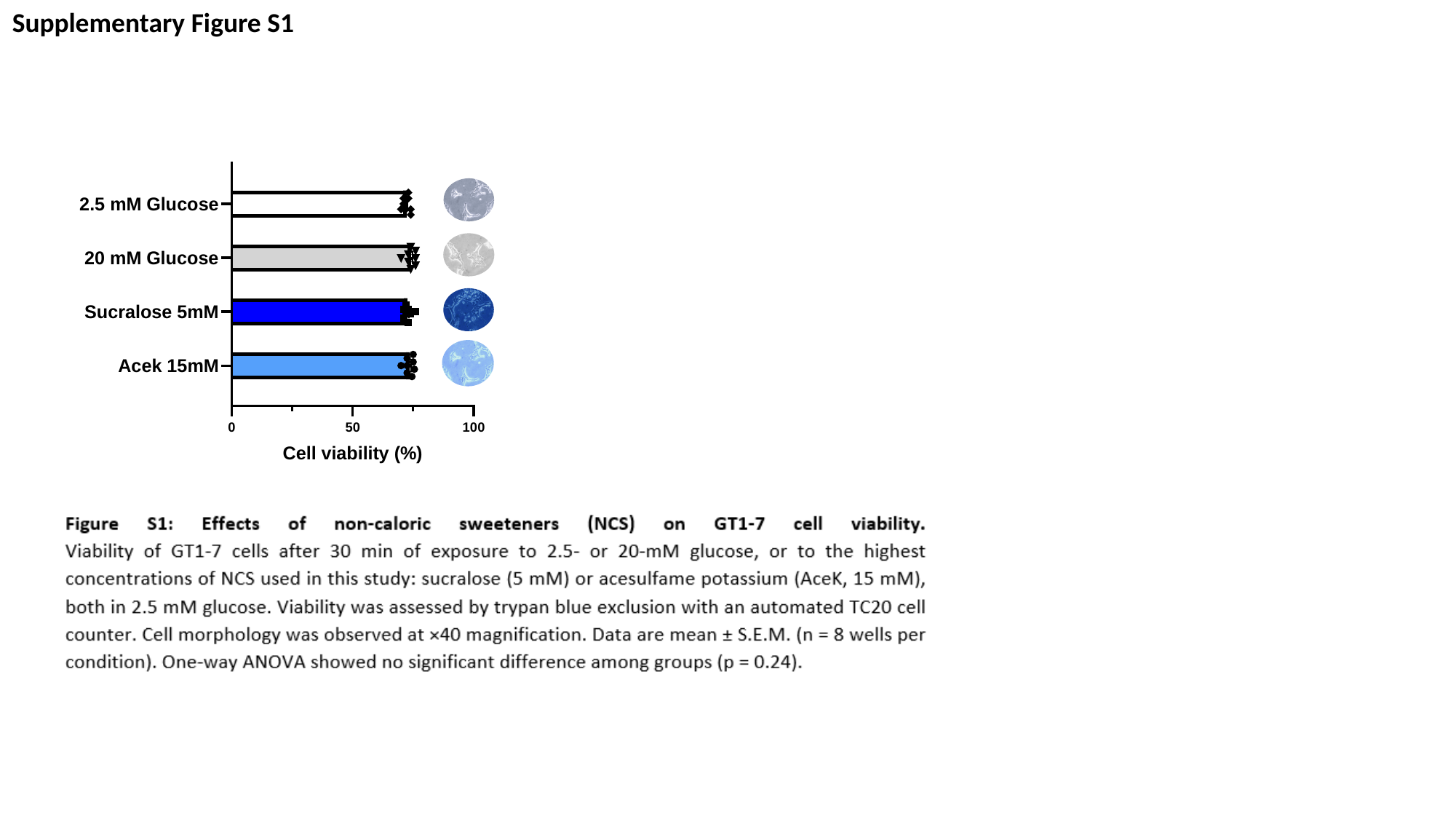

Supplementary Figure S1

### Supplemental Figure S2

#### Slide 1
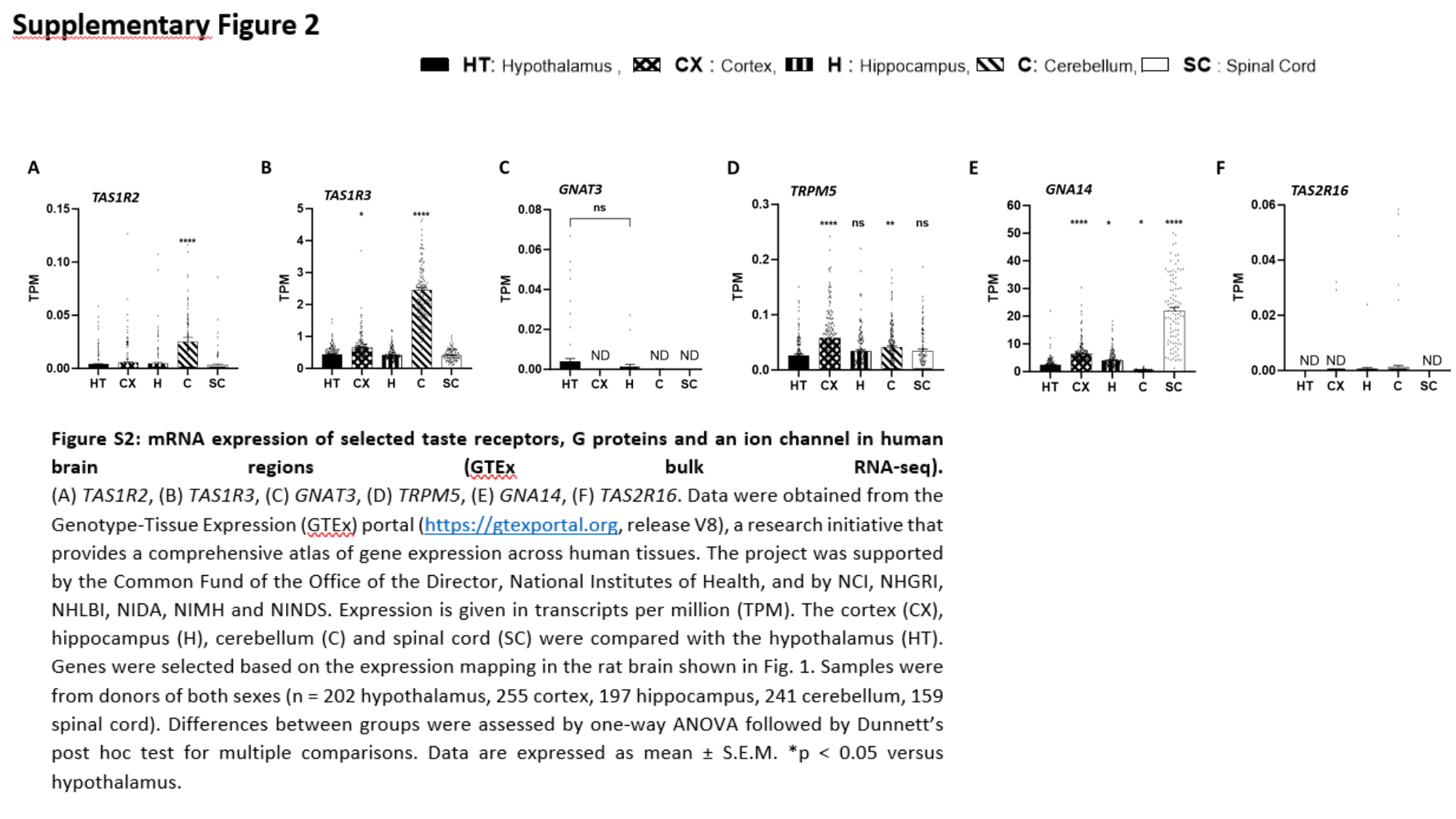
