## Supplementary Table S1 TaqMan assays for "Non-caloric sweeteners combined with glucose affect hypothalamic glucose sensing-induced insulin secretion and refeeding through neuronal cellular metabolism: *in vivo* and *in vitro* approaches"

**Supplementary Table S1. TaqMan assays used for RT-qPCR in rat and mouse samples.**

| **Species** | **Protein** | **Gene symbol** | **Assay ID** | **Use** |
| --- | --- | --- | --- | --- |
| Rat | T1R3 | *Tas1r3* | Rn00590759_g1 | Target |
| Rat | T1R2 | *Tas1r2* | Rn01515491_m1 | Target |
| Rat | Gα-gustducin | *Gnat3* | Rn00597619_m1 | Target |
| Rat | T2R116 | *Tas2r116* | Rn01462692_s1 | Target |
| Rat | Gα14 | *Gna14* | Rn01760288_m1 | Target |
| Rat | TRPM5 | *Trpm5* | Rn01479552_m1 | Target |
| Rat | PPIA | *Ppia* | Rn00690933_m1 | Reference |
| Rat | PGK1 | *Pgk1* | Rn01474008_gH | Reference |
| Mouse | T1R3 | *Tas1r3* | Mm00473459_g1 | Target |
| Mouse | T1R2 | *Tas1r2* | Mm00499716_m1 | Target |
| Mouse | Gα-gustducin | *Gnat3* | Mm01165313_m1 | Target |
| Mouse | UBC | *Ubc* | Mm01198158_m1 | Reference |
| Mouse | TBP | *Tbp* | Mm01277042_m1 | Reference |

All assays are TaqMan Gene Expression Assays (Applied Biosystems). Target genes were normalized to *Ppia* and *Pgk1* (rat samples) or *Ubc* and *Tbp* (mouse samples).
