## Supplementary appendix Gurmarin for "Non-caloric sweeteners combined with glucose affect hypothalamic glucose sensing-induced insulin secretion and refeeding through neuronal cellular metabolism: *in vivo* and *in vitro* approaches"

**Supplementary Appendix. Gurmarin used as a sweet taste receptor inhibitor**

| **Product** | Recombinant Q1-gurmarin |
| --- | --- |
| **Source** | Provided by L. Briand (Centre des Sciences du Goût et de l’Alimentation, Dijon, France); produced and purified as described in Sigoillot et al., 2012 |
| **Form** | 1.0 mg/mL in 50 mM sodium phosphate buffer, pH 7.5, sterile-filtered (0.2 µm) |
| **Molecular mass** | 4,224.0 Da (1 mg/mL = 236.7 µM) |
| **Storage** | Aliquoted, −20°C |
| **Concentration used in this study** | 10 µg/mL (≈ 2.4 µM) in the ROS assay [CONFIRM for ATP assay] |

**Description**

Gurmarin is a 35-amino-acid polypeptide, with an amino-terminal pyroglutamyl residue, isolated from the leaves of Gymnema sylvestre. In rodents it selectively inhibits responses to sweet substances without affecting responses to other basic taste stimuli such as NaCl, HCl and quinine. In vivo, its effect on taste nerve responses is long-lasting (> 2–3 h). Gurmarin inhibits rat TAS1R2/TAS1R3 and TAS1R1/TAS1R3 receptors in cellular assays: the recombinant Q1-gurmarin blocked saccharin-induced responses in rat T1R2/T1R3-expressing cells with a half-maximal inhibitory concentration of 0.030 ± 0.001 µg/mL (Sigoillot et al., 2012). The concentration used here (10 µg/mL) is therefore about 330 times this IC50.

**Preparation**

Gurmarin was supplied at 1.0 mg/mL, aliquoted and stored at −20°C, and thawed on ice or at room temperature before use. Repeated freeze–thaw cycles were avoided.
